# CardamomOT: a mechanistic optimal transport-based framework for gene regulatory network inference, trajectory reconstruction and generative modeling

**DOI:** 10.64898/2026.03.31.715390

**Authors:** Yann Maugé, Elias Ventre

**Affiliations:** COMPO, Inria Méditerranée, Cancer Research Center of Marseille, Inserm UMR1068, CNRS UMR7258, UM105, Aix-Marseille Université, SouthROCK Center of Excellence in Pediatric Oncology Research, 13273 Marseille, France

**Keywords:** gene regulatory network, mechanistic learning, transcriptional bursting, trajectory inference, optimal transport, generative modeling

## Abstract

A key challenge in inferring gene regulatory networks (GRNs) governing cellular processes such as differentiation and reprogramming from experimental data lies in the impossibility of directly measuring protein dynamics at the single-cell level, which prevents establishing causal relationships between regulator activity and target responses. In earlier work, we introduced CARDAMOM, an algorithm that uses temporal snapshots of scRNA-seq data to calibrate a GRN-driven mechanistic model of gene expression. However, this method had several limitations: it could only rely on the relative ordering of time points rather than their exact labels, imposed restrictive quasi-stationary assumptions on protein dynamics, and depended on multiple hyperparameters.

Here, we present CardamomOT, a new method based on the same mechanistic model that jointly reconstructs the GRN and unobserved protein trajectories from the data within a mechanistic optimal transport framework. By incorporating exact time labels and priors on protein kinetic rates from the literature, and substantially reducing the number of required hyperparameters, our approach addresses these limitations and substantially improves the accuracy and robustness of GRN calibration. We validate our framework on both *in silico* and experimental datasets, demonstrating computational scalability and consistently improved performance over state-of-the-art methods in both GRN and trajectory reconstruction on simulated datasets, and, on experimental datasets, reconstruction of cellular trajectories, velocity fields and latent protein levels that are mutually consistent, together with GRN structures consistent with known biology. We also show that these improvements make the calibrated mechanistic model suitable to be used as a generative model to generate testable predictions of cellular responses to unseen perturbations. To our knowledge, this is among the first methods to explicitly integrate mechanistic GRN inference, trajectory reconstruction, and simulation of realistic datasets into a unified framework for scRNA-seq time series analysis.

**Author summary:** Predicting gene regulatory interactions and understanding how cellular trajectories respond to perturbations are central challenges in cell biology and bioinformatics, yet they have long been addressed as separate problems. Only a few recent approaches leverage temporal single-cell RNA sequencing data to jointly infer gene regulatory networks (GRNs) and cellular trajectories, including our previous method CARDAMOM.

In this work, we introduce CardamomOT, a new framework that integrates optimal transport—a mathematical theory that has become widely used in computational biology in recent years—within a mechanistic modeling approach. This allows us to link cells across timepoints while jointly reconstructing the underlying, unobserved protein trajectories that drive gene regulation. By explicitly modeling these latent dynamics, our method substantially improves the robustness and accuracy of GRN inference.

In contrast to many black-box approaches, CardamomOT is grounded in a biologically interpretable mechanistic model of gene expression, incorporating key processes such as transcriptional bursting, protein translation, and degradation. This framework naturally accommodates prior knowledge on RNA and protein kinetic rates available in the literature and remains consistent with the statistical properties of single-cell transcriptomic data.

We show that CardamomOT accurately infers GRNs and reconstructs cellular trajectories on simulated data, and recovers biologically consistent GRNs and trajectories on experimental data. Moreover, because it provides a fully calibrated generative model, it can be used to predict cellular responses to unseen perturbations by modifying the inferred regulatory interactions.

## Introduction

Biological processes such as cellular differentiation and reprogramming are governed by time-dependent changes in gene expression. These dynamic processes can be partially observed at the single-cell level thanks to the development of single-cell RNA sequencing (scRNA-seq) technologies. Recent advances have enabled researchers to collect single-cell gene expression data from cell populations at multiple time points, revealing high and meaningful variability between individual cells [1, 2, 3, 4] that population-based approaches fail to capture due to averaging effects [5, 6].

The dynamics of gene expression in single cells are traditionally attributed to the underlying structure of gene regulatory networks (GRNs), which describe interactions between genes mediated by the production of regulatory proteins (e.g. transcription factors (TFs) and their upstream activators) and RNAs (e.g. miRNA, siRNA, etc.). Consequently, automated inference of GRNs from high-throughput data has become a central task in systems biology [7], driven by continuous improvements in experimental technologies. Methods such as GENIE3 [8] and its extension SCENIC [9] have become standard tools, and have been extended to account for temporal structure when available [10]. More broadly, it is now well established that GRN identifiability can be greatly enhanced by observing cellular trajectories at single-cell resolution [11].

In practice, however, such trajectories remain largely inaccessible at the genome-wide scale. Although emerging approaches (e.g., live-cell transcriptomics or Raman-based techniques [12, 13]) begin to enable longitudinal measurements, they currently lack the throughput, resolution, and temporal depth required for robust inference. Standard scRNA-seq protocols, by contrast, are destructive and preclude repeated measurements of the same cell.

As a result, most datasets consist of independent snapshots of cell populations at successive timepoints, treated as samples from a shared underlying dynamical process. This disconnect between the destructive nature of scRNA-seq and the need for longitudinal tracking poses a major challenge for GRN inference. To address it, optimal transport (OT)-based methods [14] have been widely used to reconstruct population-level trajectories by linking cells across timepoints [15, 16, 17], effectively approximating cell dynamics under a least-action principle in gene expression space. A wide range of methods have subsequently been developed for building a GRN from such trajectories, from simple differential expression analysis [18] to flow matching [19, 20] and mechanistic neuralODEs [21]. However, most OT-based trajectory inference methods rely on oversimplified assumptions about the geometry of gene expression space and consider a squared-Euclidean OT cost, which amounts to working in a flat space. In the entropically regularized version of OT used in the aforementioned references, this is equivalent to assuming that individual cells follow a stochastic differential equation (SDE) with constant diffusion rate. Trajectory reconstruction then effectively consists in finding the best gradient drift reproducing the observed dynamics under an initial Brownian-motion prior in gene expression space [22], reflecting a lack of mechanistic prior knowledge [23]. Recent approaches propose to go beyond this Brownian prior by introducing new OT-derived dynamical problems [24, 25, 26, 27], but the geometric constraints remain only partially mechanistic, and realistic priors often come at a much higher computational cost (see Appendix B).

A second important challenge arises from the limited availability of single-cell proteomics. Despite notable progress in recent years [28], current technologies can quantify only a subset of proteins, and key regulators such as TFs often remain inaccessible. As a result, statistical modeling typically relies on scRNA data whose synthesis by transcriptional bursts makes it inherently stochastic and highly variable [29, 30], with non-Gaussian distributions [31, 5]. The variable relationship between mRNA and protein levels, due to transcriptional and translational regulation, makes it difficult to functionally interpret mRNA variations to build a GRN, particularly in time-series scRNA-seq datasets where the kinetic rates characterizing mRNA and protein dynamics can impact the reconstructed GRN [32]. This challenge is generally ignored in the aforementioned OT-based methods, which operate in transcript space only and thus rely on a gene-only underlying model, without explicitly accounting for hidden variables such as proteins for which kinetic information is nevertheless available. Bertin et al. constitute a notable exception [21], using deep learning techniques to integrate latent variables such as additional genes and proxies for proteins. However, they do not explicitly model protein dynamics or kinetic rates, and the trajectory inference component still relies on a Brownian-motion prior without mechanistic constraints. To the best of our knowledge, the only class of methods directly addressing this challenge is based on mechanistic modeling of gene expression dynamics [33, 34, 35, 32, 36], all using multidimensional generalizations of the well-known two-state model [37].

In this paper, we present CardamomOT, which extends our previously published method CAR-DAMOM [32] and addresses both challenges simultaneously within a unified mechanistic optimal transport framework. Compared to CARDAMOM, CardamomOT introduces three key advances: (i) it explicitly reconstructs protein trajectories through an iterative OT-based procedure, abandoning the quasi-stationary approximation on protein dynamics; (ii) it directly incorporates exact time labels and prior knowledge on kinetic rates, increasing robustness by removing the need for time-dependent hyperparameters; and (iii) it produces a fully calibrated generative model of gene expression, enabling downstream analyses that go beyond GRN inference.

More precisely, CardamomOT calibrates the GRN-driven mechanistic model of gene expression introduced in [33, 32] from temporal snapshots of scRNA-seq data by jointly reconstructing hidden protein trajectories and a GRN capable of regenerating the observed dataset (Figure 1). This joint reconstruction is performed through an iterative expectation-maximization-like (EM-like) procedure alternating between protein trajectory inference — via an OT problem with a mechanistic, GRN-driven cost — and GRN calibration by regression on the reconstructed protein values, until convergence (Figure 1D). The method requires prior knowledge about kinetic rates of protein dynamics, can benefit from prior knowledge about kinetic rates of mRNA dynamics, and can be further constrained by known regulatory interactions.

**Figure 1.**
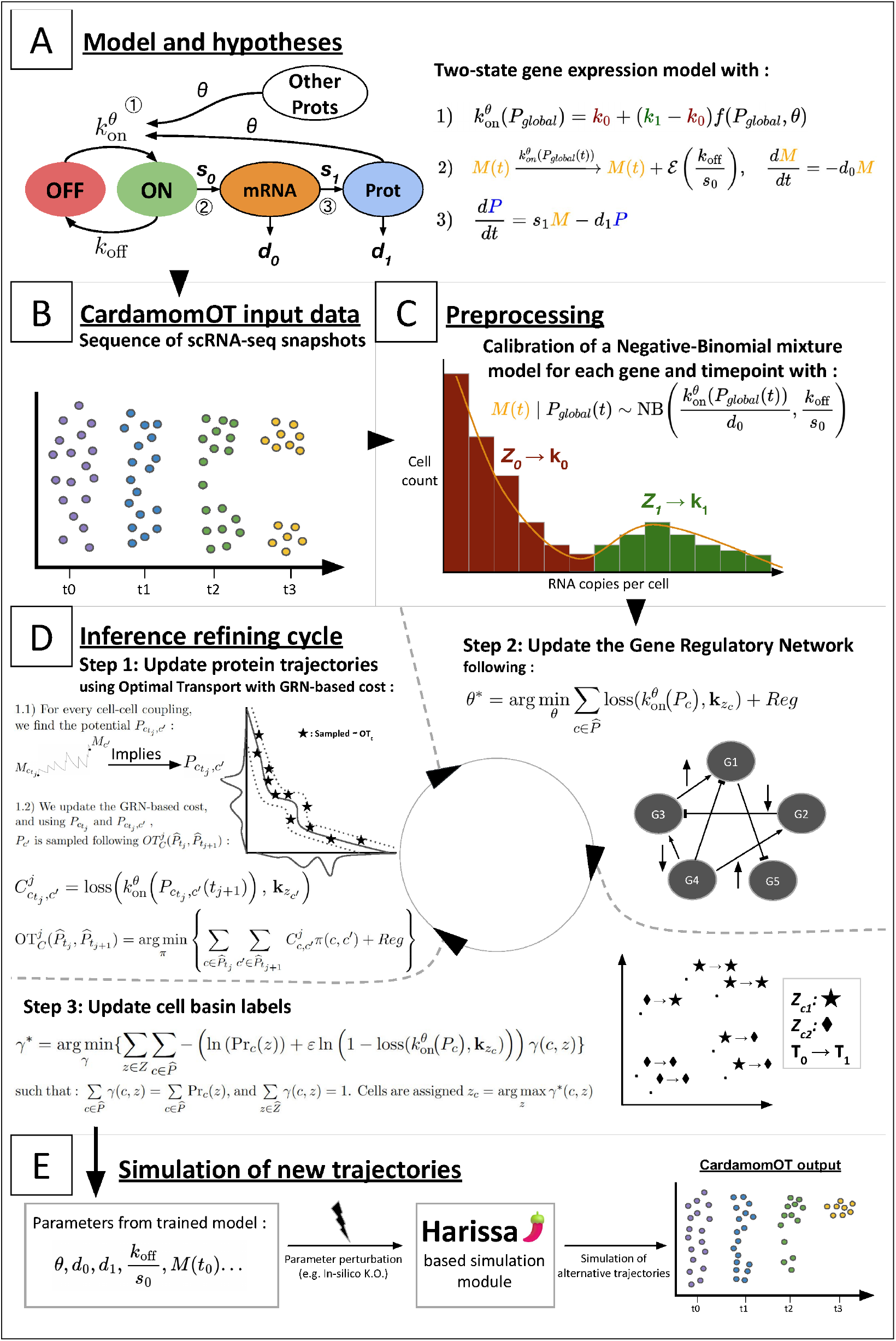
CardamomOT infers gene regulatory networks (GRNs) and protein trajectories from temporal snapshots of scRNA-seq data using a mechanistic optimal transport framework. (A) The two-state gene expression model underlying the method, in which promoter switching rates are modulated by a GRN matrix *θ*. (B) Input data consisting of independent scRNA-seq snapshots at successive timepoints. (C) Preprocessing step: a Negative Binomial mixture model is calibrated for each gene and timepoint, clustering cells into discrete promoter-activity basins. (D) Iterative EM-like inference cycle alternating between protein trajectory reconstruction via GRN-driven optimal transport (Step 1), GRN update by regression on reconstructed protein values (Step 2), and basin label refinement (Step 3). (E) The calibrated model is used as a generative module for simulating alternative trajectories and in silico perturbation experiments.

We validate CardamomOT on multiple tasks: GRN inference accuracy, reconstruction of cellular trajectories and associated velocity fields, data regeneration, and perturbation prediction. A central advantage of this approach is that the calibrated mechanistic model functions as a generative model: once trained, it can simulate new temporal snapshots that reproduce the observed data and can be used to predict the effect of unseen genetic perturbations directly *in silico* (Figure 1E). In particular, gene knockouts and overexpressions can be simulated by modifying protein levels and basal parameters in the GRN without retraining the model.

Benchmarks and applications are performed on an already published set of simulated datasets [32], an experimental dataset of sympathoadrenal differentiation, preprocessed with pseudotime in [38] from Kameneva et al. [39], and two well-known timestamped experimental datasets of differentiating cells: one characterizing the retinoic acid driven mouse embryonic stem cells differentiation, from Semrau et al. [3], and one representing the reprogramming of fibroblasts to induced pluripotent stem cells (iPSCs), from Schiebinger et al. [15]. We show, among other results, that CardamomOT correctly predicts the effect of the overexpression of Obox6 and Zfp42 on iPSC reprogramming efficiency [15] — an outcome that had been verified experimentally — using only temporal scRNA-seq data for inference and no prior knowledge of these factors’ function.

### Related works

On the GRN inference side, the mechanistic model used throughout this article was introduced together with the first method to infer a GRN from it in [33, 31], and later extended to temporal snapshots in CARDAMOM [32]. CARDAMOM, however, could not exploit experimental time labels beyond their relative ordering, relied on a quasi-stationary approximation for protein dynamics that implicitly constrained kinetic rates, did not reconstruct individual cell trajectories, and required time-dependent regularization hyperparameters that limited its robustness and prevented its use as a generative model.

On the trajectory inference side, several approaches have recently been proposed to go beyond the Brownian-motion prior. Reference Fitting (RF) [25] and GRIT [26] jointly infer cellular trajectories and GRNs using entropic OT by approximating the system dynamics as a linear Ornstein–Uhlenbeck process. This approach, however, is limited to linear systems and cannot effectively model non-linear dynamics. Wasserstein Lagrangian Flows (WLF) have been introduced to solve a dynamical OT-like problem by directly incorporating a potential energy constraint in the Lagrangian of the corresponding variational problem [24], but this comes at a high computational cost since all characteristics of the underlying process and its time-varying measure must be parametrized by neural networks. Our approach can be seen as an extension of RF and GRIT to the case where the reference process is a non-linear mechanistic process, making it suitable for modeling complex behaviors. There is also a direct link with WLF, as CardamomOT can be interpreted as solving an OT-like variational problem where the Lagrangian is induced by a simplified mechanistic process on a well-chosen latent space. These correspondences are detailed in Appendix B.

## 1 Methods

### 1.1 Mechanistic modeling of cellular dynamics

Single-cell dynamics emerge from the stochastic interplay between gene activation, mRNA synthesis, protein production, and their respective degradation processes. We adopt the hybrid two-state gene-expression model embedded in a regulatory network introduced in [33], which provides a parsimonious yet biologically grounded description of this interplay.

#### Biological model

Each gene *i* ∈ *{*1, …, *n}* has a promoter that switches stochastically between an inactive and an active state. When active, transcription fires at rate *s*_0,*i*_, producing mRNA molecules subsequently translated into proteins at rate *s*_1,*i*_; both species degrade with rates *d*_0,*i*_ and *d*_1,*i*_, respectively. For an isolated gene, the switching rates are fixed constants *k*_on,*i*_ and *k*_off,*i*_. To encode a GRN *θ* = (*θ*_*ij*_), the rates of gene *i* are replaced by protein-dependent functions, 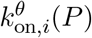, so that each entry *θ*_*ij*_ captures the sign, direction, and magnitude of the influence of gene *j* on gene *i*.

Recent experimental evidence suggests that regulation acts primarily through changes in activation frequency rather than inactivation rates [40, 41]. Accordingly, we take *k*_off,*i*_ to be independent of protein levels and focus on the *bursty* regime *k*_on,*i*_ ≪ *k*_off,*i*_, where transcription occurs in short, highly productive bursts of mean size *s*_0,*i*_*/k*_off,*i*_, which is well supported for the majority of the genes [42, 41]. Constitutively expressed genes may not satisfy *k*_on_ ≪ *k*_off_; for these, the Negative-Binomial approximation remains valid but the two-mode approximation is less appropriate. An extension to softmax activation functions which generalize the sigmoid to genes exhibiting more than two expression modes would be useful in that case.

Following [31, 32], in all applications presented in this article we adopt a sigmoid parameterization for the activation rates

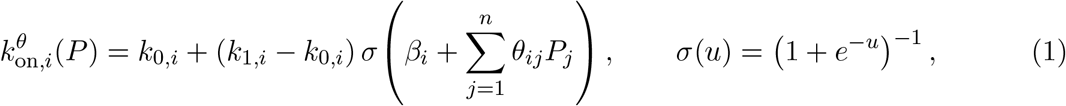

where *k*_0,*i*_ and *k*_1,*i*_ are the minimal and maximal burst frequencies of gene *i*, and *β*_*i*_ is its basal activity. We nevertheless emphasize that our method is directly applicable to a more general class of functions, that are already implemented in our Python package CardamomOT, as discussed in Section 3.

The full stochastic dynamics of mRNA *M*_*i*_(*t*) and protein *P*_*i*_(*t*) are then

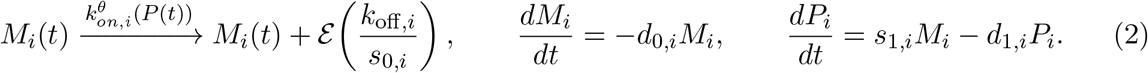

forming a hybrid stochastic-deterministic model where → denotes a stochastic jump and 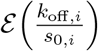 is the exponential distribution characterizing the burst size of mean *s*_0_*/k*_off_.

This model belongs to the general class of Piecewise Deterministic Markov Processes (PDMP).

#### Simplified statistical model

When bursting and mRNA degradation are fast relative to protein dynamics, the conditional distribution of *M*_*i*_(*t*) given *P* (*t*) reaches quasi-stationarity and is well approximated by a Negative Binomial (NB) distribution [29, 43, 31]:

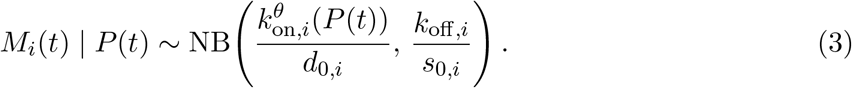

where NB(*a, b*) denotes the Gamma–Poisson mixture defined by *λ* ~ Gamma(shape *a*, rate *b*) and *M* | *λ* ~ *p* (*λ*), so that 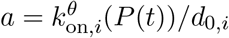 is the shape parameter (burst frequency relative to the mRNA degradation rate) and *b* = *k*_off,*i*_*/s*_0,*i*_ the rate parameter (inverse mean burst size). This yields 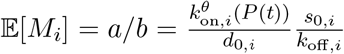 and a Fano factor 1 + *s*_0,*i*_*/k*_off,*i*_, i.e. that mean expression is the product of burst frequency and burst size, and that overdispersion is controlled by burst size. Equivalently, in the size–probability convention, NB with size *a* and probability *b/*(1 + *b*).

Assuming further that 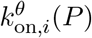 is effectively piecewise constant, taking values *{***k**_*z,i*_*}*_*z*∈*Z*_ associated with discrete promoter-activity *basins*, the marginal distribution of *M* (*t*) at population level becomes an NB mixture:

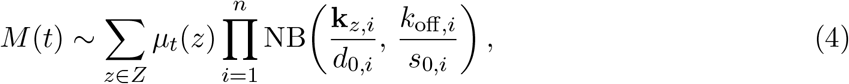

where *µ*_*t*_ is the distribution of cells among basins at time *t*. This mixture structure motivates using the basin labels *z* as a discrete, mechanistically interpretable representation of cell state.

#### Deterministic limit for proteins

In the limit where proteins evolve on a much slower timescale than mRNAs, the latter are well approximated by their conditional mean given *P*, and protein dynamics satisfy

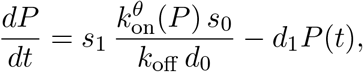

where the notation is here vectorial. Setting *s*_1_ = *d*_1_*d*_0_*k*_off_ */*(*k*_1_*s*_0_) rescales protein levels to the unit interval, yielding

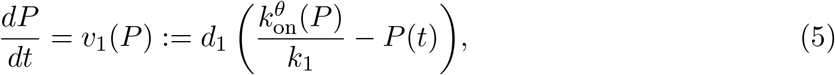

which we use both for trajectory interpolation (Appendix A) and for the postprocessing calibration of kinetic rates (Section 1.2.3).

This normalization does not alter the qualitative dynamics of the system, but simply corresponds to a rescaling of protein concentrations. Since protein levels are not directly observed, their absolute scale is not identifiable and can be fixed without loss of generality. Importantly, this choice ensures that all protein variables *P*_*i*_ lie on a comparable scale, so that the interaction coefficients *θ*_*ij*_ in Eq. (1) have a consistent interpretation in terms of regulatory strength across genes.

We emphasize that in our method presented below, we systematically adopt a *metastable* perspective (i.e., cells spend most of their time near stable attractors and undergo rare stochastic transitions between them), as characterized in [44]: the deterministic model (5) captures intra-basin dynamics, with basins corresponding to attractors of the limiting system, while stochastic fluctuations govern rare transitions between these basins in experimentally relevant regimes.

The dynamics of transitions between basins, combined with (3), define a Hidden Markov Model in which basins follow a continuous-time Markov chain and generate mRNA levels according to the GRN-dependent distribution (3). We provide further details on this perspective in Appendix B, where we refer to this model as the *phenomenological model* (see also [44]).

#### Average behavior for mRNAs

In contrast to proteins, there is no simple deterministic limit that closes the dynamics directly in mRNA space: the burst frequency 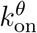 is controlled by proteins, so that *M* alone does not form a closed system. Nevertheless, given a protein state *P* (*t*), we can define an average mRNA velocity based on burst size and frequency:

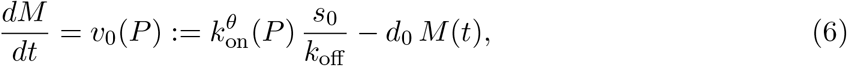

which we use for evaluation of trajectory inference in Figure S7.

We emphasize that this effective field is in general highly history-dependent, because the promoter activation regimes are driven by proteins that evolve on a slower timescale and encode the underlying regulatory logic. Any velocity model formulated directly on mRNAs alone must therefore implicitly integrate over these hidden protein states, and thus over the mechanistic structure of the GRN, leading to an effective regulation that is no longer explicitly mechanistic [21]. CardamomOT keeps the dependence on *P* explicit and reconstructs protein trajectories alongside the GRN, so that both protein and mRNA dynamics remain grounded in the same mechanistic model.

### 1.2 Overview of CardamomOT

CardamomOT calibrates the mechanistic model above as a *generative model* for scRNA-seq time series. Starting from raw count matrices at successive timepoints *t*_1_ *< …< t*_*N*_, it jointly infers the GRN *θ*^*^ and hidden protein trajectories 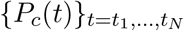 for each cell *c*, then uses the calibrated model to simulate new data or predict perturbation effects. The method requires prior knowledge of protein degradation rates *d*_1_, and can optionally incorporate mRNA degradation rates *d*_0_ and known regulatory interactions *θ*^0^.

#### 1.2.1 Preprocessing: inferring the NB mixture and initializing basin labels

We first calibrate the NB mixture model (4) gene by gene across timepoints, following [32]. For each gene *i* and timepoint *t*_*j*_, we fit NB(*a*_*j,i*_, *b*_*i*_) with *b*_*i*_ shared across time, and set

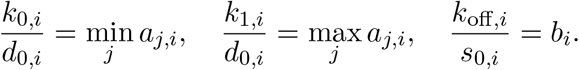

This yields, for each cell *c*, a probability matrix Pr_*c*_ of size |*Z*| *× n* encoding the likelihood of belonging to each basin. Cells are initialized by maximum-likelihood assignment:

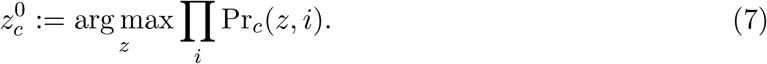

If a prior GRN *θ*^0^ is available, this assignment can already incorporate network information via the GRN-constrained update described in Step 3 below.

#### 1.2.2 Iterative inference of GRN and protein trajectories

The core novelty of CardamomOT is to abandon the quasi-stationary approximation 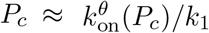 used in CARDAMOM, and instead reconstruct protein trajectories 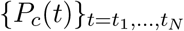 through explicit trajectory inference. Since protein dynamics are driven by the GRN, trajectory inference and GRN reconstruction are tightly coupled: the GRN shapes the interpolation cost, and the reconstructed trajectories in turn constrain the GRN. We therefore alternate between the two in an EM-like loop.

##### Step 1: Protein trajectory reconstruction via mechanistic OT

Given a current GRN *θ* and basin labels *{z*_*c*_*}*, we seek, for each cell 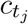 at time *t*_*j*_, its most likely descendant 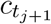 at *t*_*j*+1_ (i.e., the cell at the next timepoint that most plausibly derives from the same lineage). For each candidate cell *c*′ observed at *t*_*j*+1_, we compute a candidate protein vector 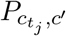 by integrating the deterministic ODE (5) from 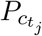 towards a state consistent with *c*′; this interpolation accounts for a possible single mode switch along the interval (see Appendix A for details). The cost of the candidate is

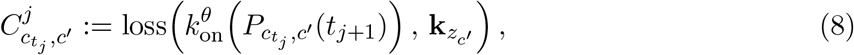

where 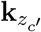 denotes the target burst-mode vector associated with the basin of *c*′, and loss is typically taken as the cross-entropy, consistent with the sigmoid parametrization of 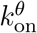 in Eq. (1).

We then solve the entropically regularized OT problem

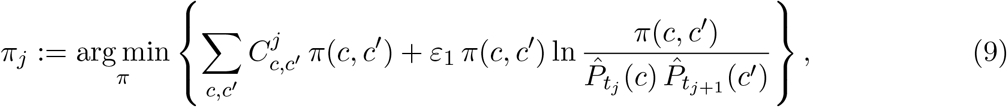

subject to the marginal constraints 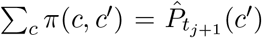 and 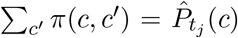. Here 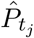 denotes the empirical distribution of cells at time *t*_*j*_, and *ε*_1_ plays the role of a temperature parameter, typically decreased linearly over iterations from 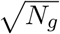 to 0.01, where *N*_*g*_ is the number of genes.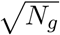 is chosen to counterbalance the typical scale of the cost matrix (8): at the first iterations, the number of incorrectly labeled genes scales as the square root of the total number of genes.

The optimal coupling *π*_*j*_ is then used to sample a descendant *c*′ for each ancestor 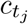, and we set 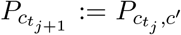. Repeating this from *t*_1_ to *t*_*N*−1_ yields full protein trajectories for all sampled cells.

##### Step 2: GRN update

Given the reconstructed protein trajectories, we update the GRN by minimizing the loss between GRN-predicted burst rates and mixture-derived basin modes over all cells and timepoints:

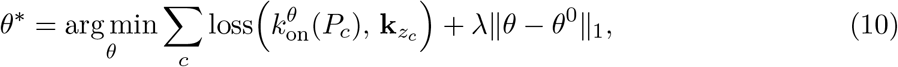

where 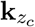 is the burst-mode vector associated with the basin of cell *c*. Compared to CAR-DAMOM, all timepoints now contribute jointly through their reconstructed protein values, and a simple *l*1 penalty with respect to the prior GRN *θ*^0^ replaces the custom time-dependent regularization. The regularization strength *λ* thus controls the sparsity of the inferred network. We set it as 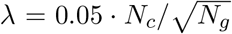, where *N*_*c*_ is the number of cells, reflecting the heuristic that the number of active regulators per gene scales at most as 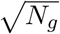, and that a per-gene basin label attribution error of 5% is a reasonable target for reliable inference.

##### Step 3: Basin label refinement

Given the updated GRN *θ*^*^, basin labels are refined by solving a GRN-constrained assignment problem that balances the per-cell NB log-likelihood with consistency between inferred proteins and candidate basin modes:

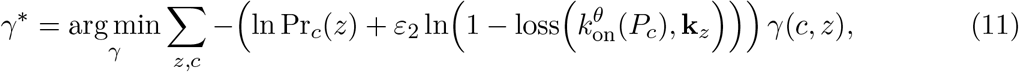

subject to Σ_*c*_ *ϒ* (*c, z*) = Σ_*c*_ Pr_*c*_(*z*) and Σ_*z*_ *ϒ* (*c, z*) = 1. Here, the quantity loss 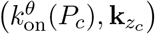 is renormalized to be in [0, 1], such that the ln is well defined. The parameter *ε*_2_ controls how strongly consistency with the mechanistic model influences label assignment; it is typically increased linearly over iterations from 0 to 0.5, so that it never completely overwhelms the NB likelihood.

Cells are then reassigned as *z*_*c*_ = arg max_*z*_ ϒ^*^(*c, z*).

##### Algorithm summary

Steps 1–3 are iterated until convergence, typically within 20–40 iterations:

###### Algorithm 1

CardamomOT

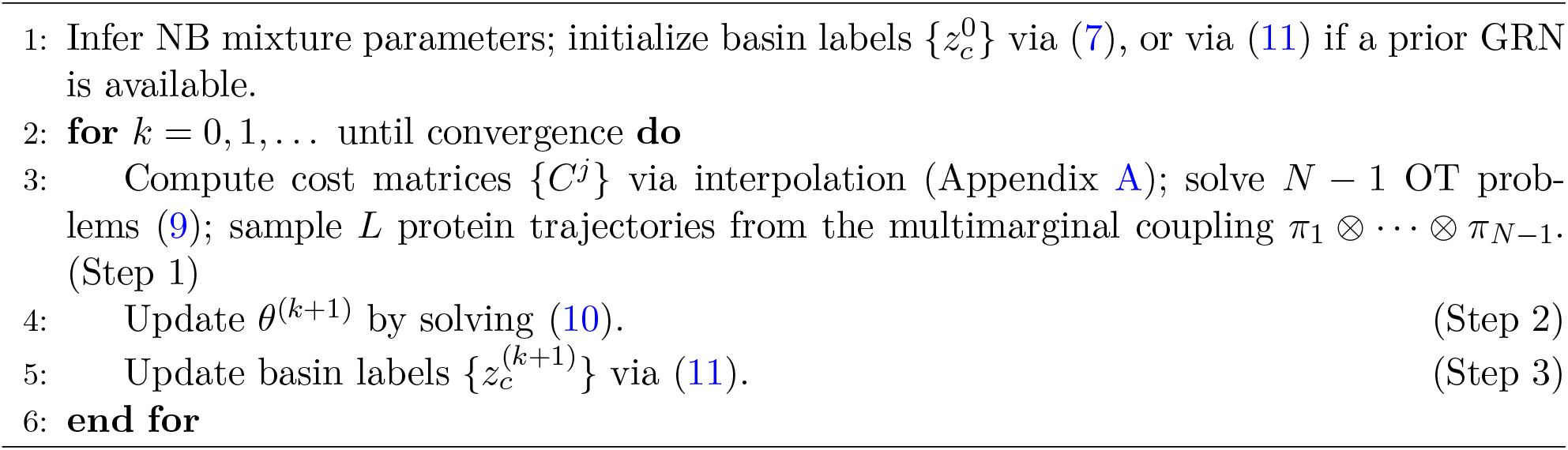

#### 1.2.3 Postprocessing: kinetic rate recalibration via differentiable ODE

Once the GRN *θ*^*^ and protein trajectories *{P*^*l*^(*t*)*}*_*l*=1,…,*L*_ have converged, we recalibrate kinetic rates so that simulations from the model reproduce the observed dynamics. This step can be interpreted as fitting the parameters of an SDE approximating the protein dynamics in the metastable regime, theoretically derived from the mechanistic model in [44].

In practice, we use a two-step approximation.

##### Step (i): protein degradation rate *d*_1_

Rather than a generic black-box neural network, the ODE used here directly instantiates the mechanistic vector field of Eq. (5), 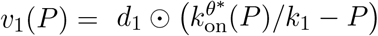, with the converged GRN *θ*^*^ and *k*_1_ held fixed. Only the protein degradation rates *d*_1_ ∈ ℝ^*n*^ are trainable, reparametrized through a softplus to enforce positivity; we additionally allow a per-gene multiplicative correction on the sigmoid argument of 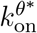, initialized at 1 and regularized toward 1 (see below), to absorb residual scale mismatches left over from the protein normalization discussed after Eq. (5).

For each reconstructed trajectory *l* and each pair of consecutive observed timepoints (*t*_*j*_, *t*_*j*+1_), the ODE is integrated forward with the differentiable ODE solver of <monospace>torchdiffeq</monospace>, starting from the *reconstructed* protein state *P*^*l*^(*t*_*j*_) rather than from the model’s own prediction at the previous interval. This *teacher-forcing* scheme prevents integration error from compounding across the many intervals of a full time series and is now standard practice for calibrating dynamical models on single-cell time series, as in PRESCIENT [45] and STORIES [46].

The training loss is the mean squared error between the integrated endpoint 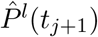 and the reconstructed observation *P*^*l*^(*t*_*j*+1_), summed over all trajectories and consecutive timepoint pairs,

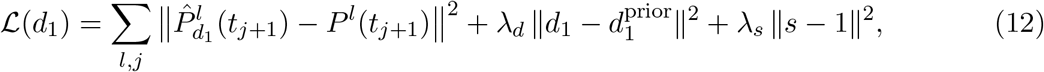

where *s* denotes the per-gene scale correction described above, and *λ*_*d*_, *λ*_*s*_ = 0.01, 1, respectively, shrink the fitted rates toward the literature-informed prior 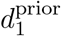 and the scale toward its neutral value, respectively. Gradients are backpropagated through the ODE solver and *d*_1_ (and *s*) are updated with Adam until convergence of the loss (Algorithm 2).

##### Step (ii): mRNA degradation rate *d*_0_

We calibrate the mRNA degradation rate *d*_0_ by fitting the variance of stochastic fluctuations around the deterministic trajectories, so that simulated expression noise matches the observed variability. Concretely, in the bursty regime, the protein increment over a short interval *dt* is driven by a compound Poisson process with burst rate 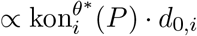 and mean burst size ∝ *d*_1,*i*_*/*(*k*_1,*i*_ *d*_0,*i*_) [35]. By Campbell’s theorem, this predicts, at leading order,

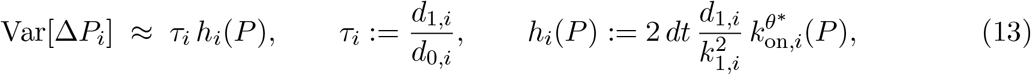

so that *τ*_*i*_ can be estimated by matching the squared residuals of the Step (i) ODE predictions (used as a proxy for Var[Δ*P*_*i*_]) against *h*_*i*_ evaluated along the reconstructed trajectories. We use the regularized, closed-form method-of-moments estimator

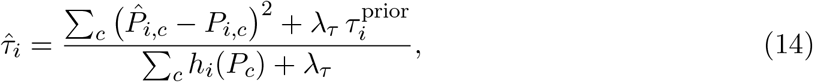

pooling over cells *c* and, optionally, over timepoints; *λ*_*τ*_ = 1 shrinks 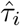 toward a literature-informed prior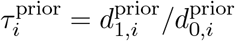 (by default corresponding to an mRNA half-life of 9 h and a protein half-life of 46 h, i.e. 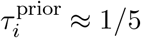, consistent with the adiabatic regime *d*_1_ ≪ *d*_0_ required by the model), unless gene-specific prior degradation rates are supplied by the user, and *d*_0,*i*_ is recovered as 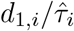.

These refined kinetic parameters are thus built such that the model generates trajectories with realistic timescales and levels of stochasticity.

###### Algorithm 2

ODE recalibration of protein degradation rates *d*_1_ (Step (i))

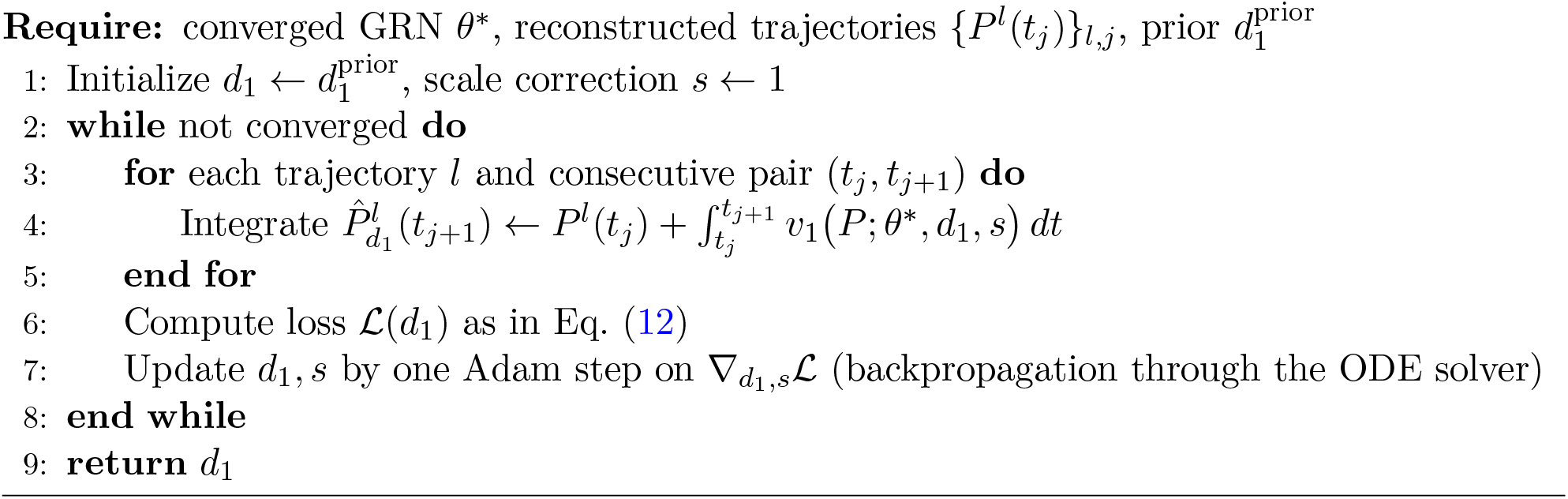

Overall, the method outputs: the inferred GRN *θ*^*^ (signed and directed); the reconstructed protein trajectories with basin labels and burst-mode vectors; and all parameters of the mechanistic model described in Section 1.1, together forming a calibrated generative model ready for simulation and *in silico* perturbation experiments.

## 2 Results

We validated the CardamomOT pipeline on both synthetic and experimental datasets, demonstrating its ability to jointly infer GRNs and cellular trajectories from temporal snapshots of scRNA-seq data, and to generate realistic simulated datasets. The method was tested on five benchmark network topologies, previously introduced in [32], and three experimental datasets: (i) mouse embryonic stem cell (mESC) differentiation [3] (9 timepoints, ~ 1,300 cells), (ii) sympathoadrenal differentiation ordered by pseudotime [38] (6 pseudo-timepoints, ~ 1,800 cells), and (iii) mouse embryonic fibroblast (MEF) reprogramming to induced pluripotent stem cells (iPSCs) [15] (35 timepoints, ~ 250,000 cells). In what follows, we present results that illustrate the complete CardamomOT workflow, from preprocessing to perturbation analysis, and show how the calibrated mechanistic model can be used as a comprehensive tool for dynamic single-cell analysis.

### 2.1 Simulation of temporal snapshots

To evaluate CardamomOT’s performance, we first generated synthetic datasets using the HARISSA package [47], which simulates biologically stochastic trajectories following the same mechanistic model as CardamomOT, described in Section 1.1. We used five benchmark network topologies from [32]: (i) FN4, a 4-gene feedforward network; (ii) FN8, an 8-gene feedforward network; (iii) BN8, an 8-gene bifurcating network; (iv) CN5, a 5-gene cascade network; and (v) a tree topology with varying numbers of genes (5, 10, 20, 50, 100 genes).

For each network, we simulated *in vitro*-like perturbation experiments by first equilibrating the system for *t <* 0, then introducing a virtual stimulus gene fixed at maximal protein level at *t* = 0, triggering a transition toward a new stochastic steady state. Temporal snapshots were generated by independently sampling cells at timepoints equally distributed from *t* = 0 to *t* = 96 hours. The model parameters (*k*_0,*i*_, *k*_1,*i*_, *s*_0,*i*_*/k*_off,*i*_, *β*_*i*_, *θ*_*ij*_) and degradation rates (*d*_0,*i*_, *d*_1,*i*_) for all genes used for these benchmarks are available online with the code repository.

This simulation framework allowed us to evaluate CardamomOT’s performance under controlled model-matched conditions where the ground truth GRN and protein levels are known, as well as true cellular trajectories when simulating without killing cells at each timepoint. This establishes best-case performance when model assumptions are satisfied, and all competing methods are evaluated under identical conditions. We then apply the method to experimental datasets, where validation is more challenging.

### 2.2 The statistical mixture model captures both mechanistic parameters and biological cell states

The first step of CardamomOT (preprocessing) consists in calibrating the NB mixture model described in Equation (4) for each gene and timepoint (Figure 1C). This step serves two critical purposes: (i) inferring the mechanistic parameters (*k*_0,*i*_, *k*_1,*i*_, *k*_off,*i*_*/s*_0,*i*_) from the data, and (ii) clustering cells into discrete promoter-activity basins that reflect their underlying regulatory states.

We first validated this approach on the simulated datasets, where ground truth parameters are known (Table S1). The burst size parameter *k*_off,*i*_*/s*_0,*i*_ is recovered accurately across all benchmark networks (0.017–0.019 against a reference value of 0.02), and the minimal burst frequency *k*_0,*i*_*/d*_0,*i*_ is estimated close to zero. The maximal burst frequency *k*_1,*i*_*/d*_0,*i*_ is systematically underestimated by ~20–30%, which follows from the estimation procedure rather than from a poor fit: *k*_1,*i*_ is the saturating value of the sigmoid in Eq. (1), whereas max_*j*_ *a*_*j,i*_ estimates the largest burst frequency actually attained in the data, and regulatory inputs do not fully saturate 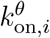 in these simulations.

In addition, CardamomOT precisely classifies cells into basins using maximum likelihood for each gene, with an average of 85%–93% correct classification across the datasets (Table S1).

For experimental datasets, where ground truth is unavailable, we assessed the quality of the statistical model fit through multiple metrics:

- **Low-dimensional embeddings:** We performed UMAP dimensionality reduction on both the original scRNA-seq data and data simulated from the calibrated statistical model. The simulated data preserved the overall temporal structure of the experimental data in UMAP space (Figure 2A,B), indicating that the statistical model captures the essential features of the gene expression landscape. In all comparisons using UMAPs throughout the paper, a low-dimensional embedding is learned from the concatenated dataset and the coordinates are then used to plot each dataset separately.
- **Marginal distributions:** The inferred NB mixture models are consistent with the observed marginal distributions and temporal patterns of gene expression for all timepoints, as illustrated in Figure 2E–H for selected genes exhibiting strong connectivity in the inferred GRN of each dataset (Figures S1–S3).
- **Correlations:** We assessed the ability of the NB mixture model to reproduce pairwise gene expression correlations observed in the reference data (Figure S5). The inferred model captures the overall correlation structure, with gene pair correlations in the NB mixture closely tracking those in the reference data across all three datasets. However, the strongest correlations tend to be slightly underestimated. This is consistent with a known limitation of the NB mixture framework: by assuming conditional independence of genes given the basin label, the model cannot capture residual correlations between genes that remain correlated within individual basins. As a result, intra-basin co-regulation between highly correlated genes is partially lost, leading to a mild attenuation of the highest correlation values.
- **Cell type identification:** We trained Random Forest classifiers on the experimental data using cell type annotations (when available) or clustering-based labels, and evaluated their performance on data simulated from the calibrated statistical model. The classifiers achieved high accuracy on reconstructing the relative proportions of cell types (Figure 2C,D), demonstrating that the inferred basin labels correspond to biologically meaningful cell states. This evaluation strategy, based on cross-dataset classification, has become standard in single-cell analysis [48] and provides an objective measure of how well the statistical model captures cell type-specific expression patterns. These results are particularly striking for the Kameneva and Schiebinger datasets, which exhibit non-trivial branching dynamics that are shown to be well distinguished by the statistical model.

**Figure 2.**
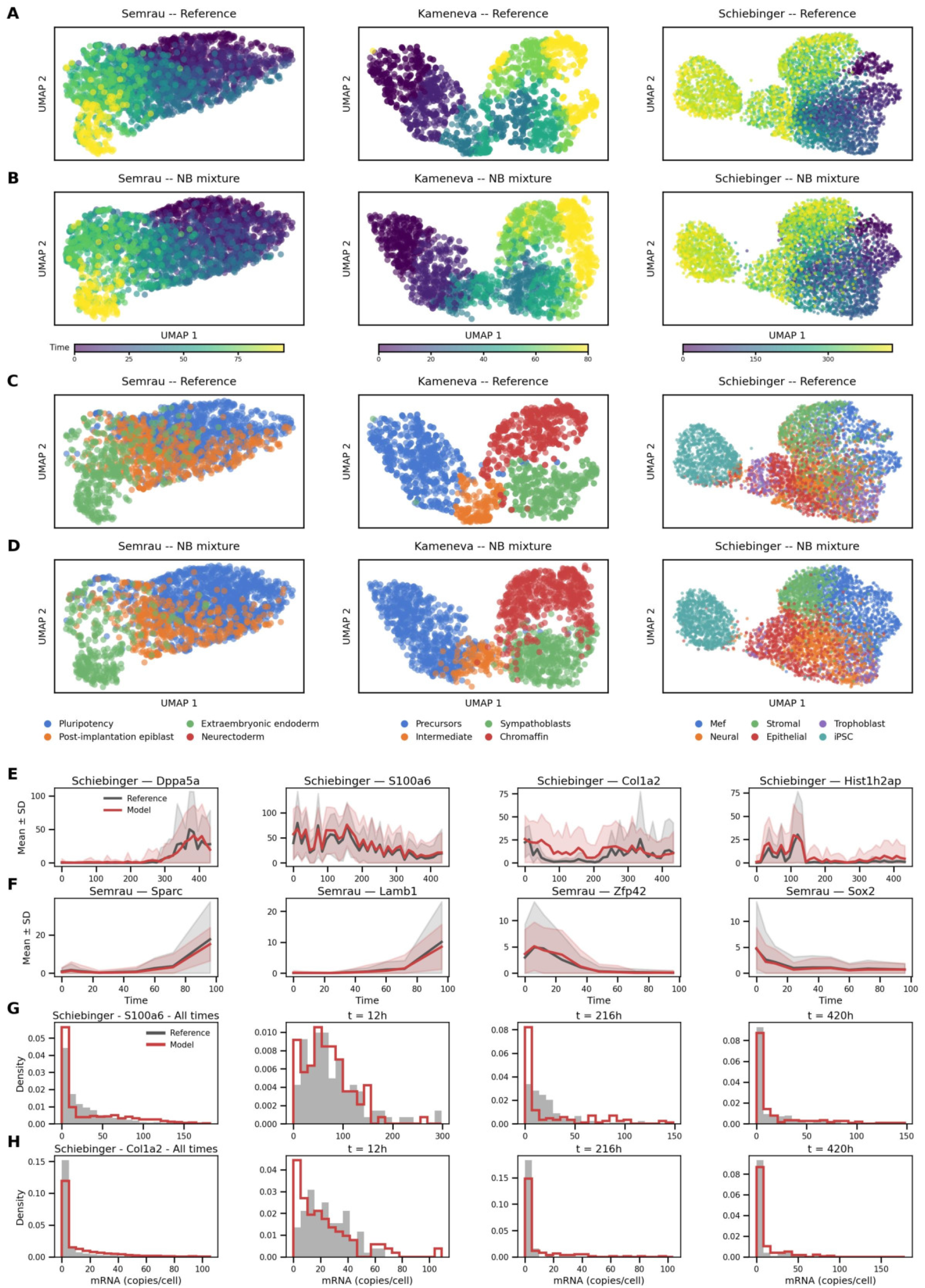
(A–D) UMAP embeddings of reference scRNA-seq data (A, C) and data simulated from the CardamomOT NB mixture model (B, D), colored by timepoint (A–B) and cell type (C–D), for the Semrau, Kameneva and Schiebinger datasets. (E–F) Temporal profiles of mean *±* SD gene expression across all cells for selected key regulators, comparing reference data (gray) and model simulations (red), for the Schiebinger (E) and Semrau (F) datasets. (G–H) Marginal mRNA distributions at selected timepoints for S100a6 (G) and Col1a2 (H) in the Schiebinger dataset, showing close agreement between model and reference data across all timepoints.

Taken together, these results show that the preprocessing step of CardamomOT not only infers mechanistically interpretable parameters but also produces a discrete representation of cellular states (basin labels) that forms the foundation for the subsequent joint GRN and trajectory inference.

### 2.3 Benchmarking CardamomOT: performance, robustness and scalability of GRN inference

We next evaluated CardamomOT’s ability to infer GRN structure from temporal snapshots. We compared CardamomOT against several state-of-the-art methods: (i) CARDAMOM [32], the previous version of our method; (ii) RF [25], a recent OT-based method that jointly infers trajectories and linear GRNs; (iii) GENIE3 [8], a widely used random forest-based method; and (iv) SINCERITIES [49], a method designed for temporal scRNA-seq data. For methods that infer undirected edges, we additionally compared against PIDC [50] and Pearson correlation, used as a sanity check to ensure that good performance is not solely due to highly correlated data that would be easy to detect.

All methods were evaluated using the Area Under the Precision-Recall curve (AUPR), a standard metric for GRN inference that is robust to class imbalance (i.e., the fact that most gene pairs are not connected). Results were averaged over 10 independent simulations for each network topology.

#### CardamomOT achieves scalable GRN inference across network topologies on model-matched simulated data

On all benchmark networks, CardamomOT achieved consistently higher AUPR scores than all competing methods (Figure 3A). The improvement was particularly pronounced for the more complex networks (FN8, BN8, and trees with more than 10 genes), where CardamomOT’s mechanistic modeling and trajectory inference provided a substantial advantage over methods that rely solely on correlation-based or quasi-static approximations. CardamomOT infers both the sign and direction of regulatory interactions, which is crucial for understanding causal relationships in GRNs. When comparing against methods that only infer undirected edges (Figure 3B), CardamomOT maintained its performance advantage, highlighting the value of mechanistic modeling for resolving regulatory directionality.

**Figure 3.**
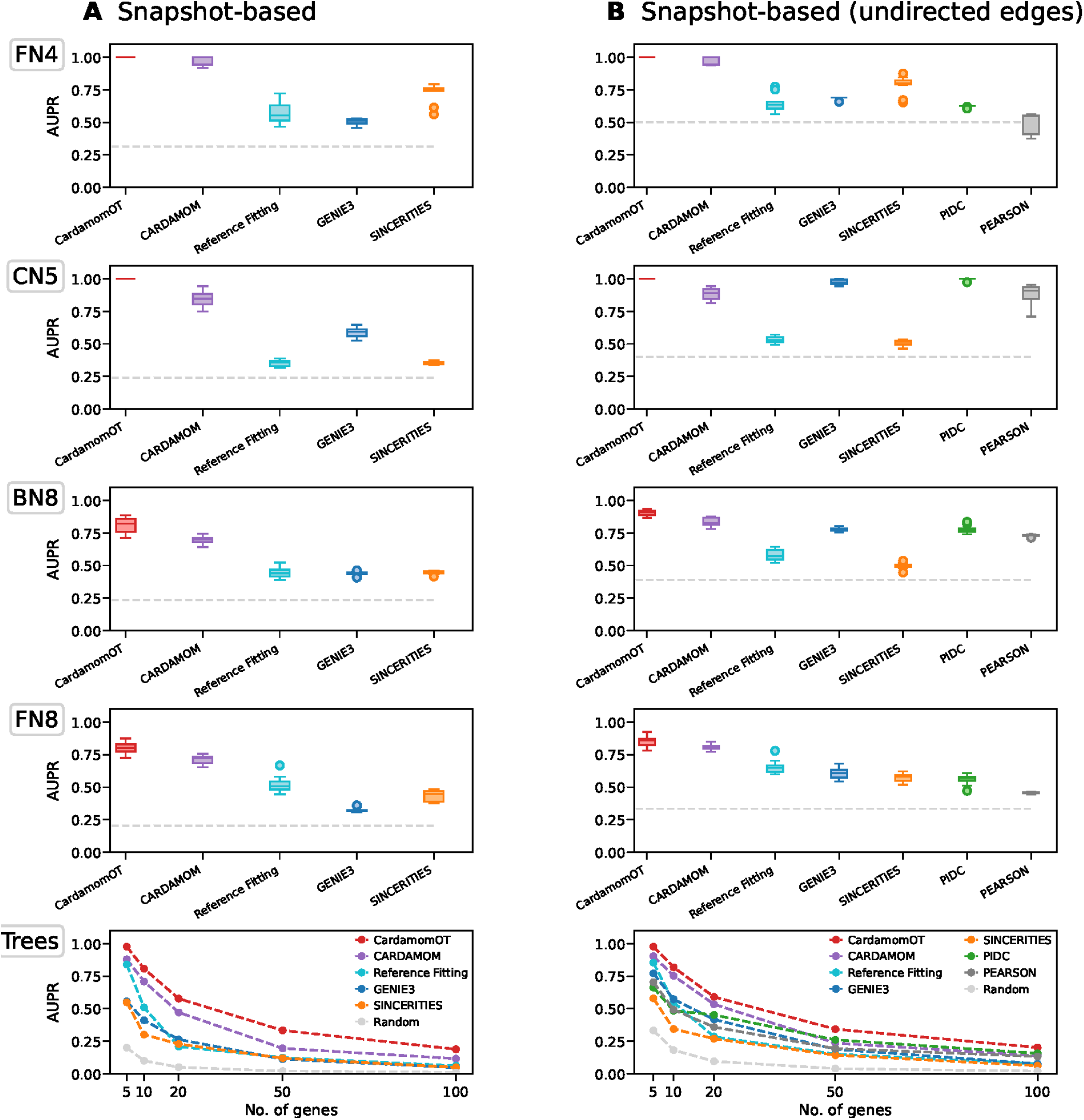
CardamomOT outperforms competing methods on GRN inference benchmarks. Performance is measured by the Area Under the Precision-Recall curve (AUPR), averaged over 10 independent simulations per network. The dashed gray line indicates the random baseline. (A) Directed GRN inference from temporal snapshots on four benchmark network topologies (FN4, CN5, BN8, FN8) and tree networks of increasing size (5–100 genes). (B) Same comparison for undirected edge inference, including additional correlation-based methods (PIDC, Pearson). Boxplots throughout the figure show median, interquartile range (box) and 1.5x interquartile range (bars).

The convergence of the inference procedure is illustrated in Figure S8: the training loss decreases monotonically and stabilizes within 20–25 optimization steps across all benchmark networks, while the AUPR score rises rapidly in the first iterations and plateaus at its final value. This indicates that the iterative EM-like procedure converges reliably and that the reported AUPR scores correspond to fully converged models rather than intermediate states.

Moreover, the runtimes reported in Table S2 show that CardamomOT scales well to realistic dataset sizes: even for networks of 100 genes with 1,000 cells, inference completes in under one minute, and CardamomOT remains faster than GENIE3, a widely used reference for GRN inference. The true limiting factor for scaling to larger networks is therefore not computational cost but the identifiability of the network structure: with *n* genes, the number of GRN parameters grows as *n*^2^, and the information available in temporal snapshots becomes insufficient to robustly constrain all interactions beyond a certain network size. This identifiability ceiling is further discussed in Section 3.

#### Iterative refinement significantly improves GRN inference accuracy

To assess the contribution of the iterative refinement procedure (Steps 1–3 in Section 1.2.2), we compared CardamomOT against several ablated versions: (i) a version that directly returns the network and trajectories inferred after the first round of the iterative procedure, i.e. the network computed from the trajectories associated with an OT coupling between basins; (ii) a version with random initialization of the OT coupling instead of the OT-based initialization; and (iii) a version combining both modifications. The full CardamomOT method consistently outperformed ablated versions (i) and (iii) (Figure 4A–E), demonstrating that the iterative refinement procedure does contribute to improved inference accuracy. Remarkably, the iterative procedure converged to similar GRN structures regardless of initialization (compare “CardamomOT” and “random init” in Figure 4A–E), suggesting that the benchmark networks exhibit sufficient identifiability for the method to escape local optima.

**Figure 4.**
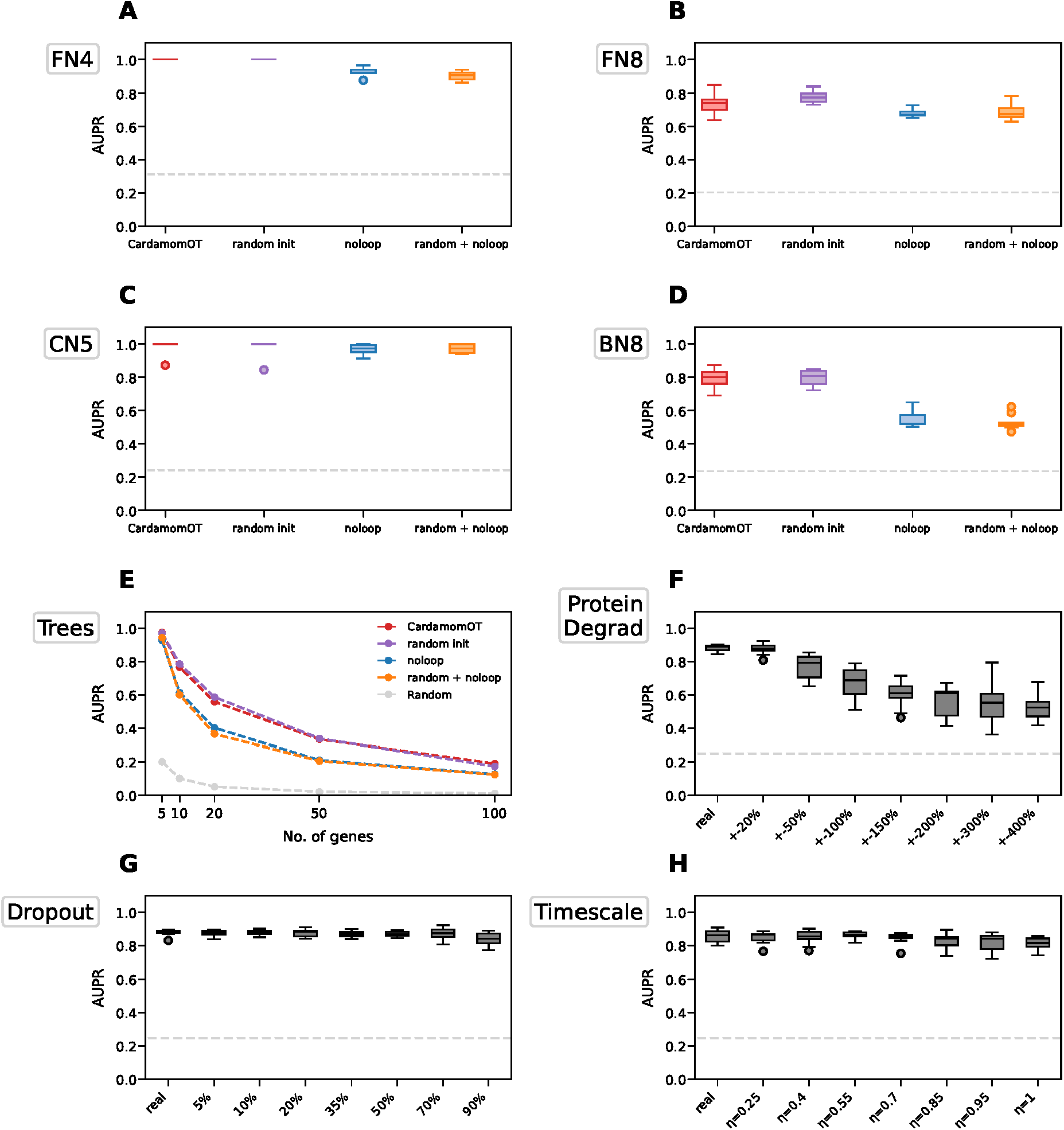
(A–E) Directed GRN inference performance (AUPR) averaged over 10 independent simulations per network comparing CardamomOT against three ablated variants: random OT initialization (random init), removal of the loop procedure (noloop), and their combination (random + noloop), on four benchmark networks and tree topologies of increasing size. (F) Robustness to uncertainty in protein degradation rates *d*_1_: AUPR averaged across 10 independent simulations for all four benchmark networks when input degradation rates are perturbed from ±20% to ±400% of their true values, compared to the performance obtained with the real rates. (G) Robustness to dropout in input data: AUPR averaged across 10 independent simulations for all four benchmark networks for base data and 7 increasing levels of binomial dropout rangin from 0% (no dropout, renamed real in the figure) to 90%. (H) Robustness in variability in timescales: AUPR averaged across 10 independent simulations for all four benchmark networks for base data and 7 increasing levels of variability in timepoint sampling. For each replicate, inter-timepoint gaps were drawn from Dirichlet(*α* = 1*/η*^2^) with endpoints fixed, where *η* ranges from 0 (equispaced timepoints, renamed real in the figure) to 1 (uniformly sampled timepoints). Boxplots throughout the figure show median, interquartile range (box) and 1.5x interquartile range (bars).

#### CardamomOT is robust with respect to prior estimation of kinetic rates

A critical advantage of CardamomOT over CARDAMOM is that it directly incorporates protein degradation rates *d*_1_ into the inference procedure (see Equation (5) and Step 1 of Section 1.2.2), allowing a much better approximation of protein trajectories than the quasi-static approximation of the previous method. Protein half-lives in mammalian cells typically range 40–100 h, with a broader range of roughly 10–200 h in more extreme cases [51, 52]. Uncertainty on these estimates can be substantial: Rolfs et al. [53] report a median uncertainty of *±*0.9 days (≈22 h) across a mouse tissue atlas of 3106 proteins. For a protein with a half-life in the 40–50 h range this corresponds to roughly *±*50% relative uncertainty, and to a comparable or larger perturbation of the degradation rate *d*_1_ = ln 2*/t*_1*/*2_, since the two are inversely related. To evaluate robustness to uncertainty in these rates, we ran CardamomOT on simulated data while systematically perturbing the input degradation rates from *±*20% up to *±*400% of their true values. The method maintained high AUPR scores (averaged across FN4, FN8, BN8 and CN5) even with moderate perturbations (*±*50%), and clearly outperformed random guessing and competing benchmark methods even with very large perturbations (*±*200–400%, Figure 4F). Typical literature-derived estimates therefore fall around the upper end of the range in which CardamomOT retains high accuracy (Figure 4F), while short-lived proteins and the tail of the uncertainty distribution may exceed it — which is why we assessed robustness up to *±*400%.

#### CardamomOT is robust to technical noise

scRNAseq measurements are prone to technical noise due to inefficient capture and sequencing. To test CardamomOT robustness to technical noise, we performed inference on simulated datasets with increasing levels of binomial dropout (5% – 90%, averaged across FN4, FN8, BN8 and CN5). CardamomOT maintains high AUPR scores with slightly increasing variability at high levels of dropout, confirming robustness to moderate technical noise (Figure 4G).

#### CardamomOT is robust to temporal variability in measurements

scRNAseq experiments commonly show irregular measurement schedules. We thus tested CardamomOT robustness to variation in measurement times by performing inference on simulated datasets on a scale going from equally spaced timepoints to randomly sampled timepoints. Average AUPR across FN4, FN8, BN8 and CN5 only slightly degrades highly variable scales, confirming CardamomOT capability to work with different measurement schedules (Figure 4H).

Overall, these results establish CardamomOT as a robust method for GRN inference on the benchmark considered.

#### Interpreting the ordering of methods in Figure 3A

An important detail in Figure 3A deserves comment: CARDAMOM, which does not reconstruct explicit cellular trajectories and instead relies on a quasi-stationary approximation [32], nonetheless outperforms RF, which does perform trajectory inference via optimal transport. This ordering should not be read as a general ranking of GRN inference methods, as evidence that trajectory reconstruction is unhelpful in general, or as evidence that OT with a Brownian/linear reference is worse than skipping trajectory inference altogether. Rather, it reflects a mismatch in how well each method’s underlying dynamical assumptions fit the statistical structure of the data.

CARDAMOM’s statistical model is the Negative Binomial approximation derived from the very mechanistic model used to generate the simulated datasets (Section 2.1), so it is, in this specific sense, correctly specified for these data and can recover accurate parameter estimates without tracking cells across timepoints. RF does reconstruct trajectories, but under a linear Ornstein– Uhlenbeck reference process that neither exploits this NB structure nor captures the nonlinear, GRN-driven dynamics of the benchmark networks; this mismatch degrades both the inferred trajectories and the GRN subsequently built from them. CardamomOT is designed precisely to combine the strengths of both, by pairing explicit trajectory reconstruction with the same nonlinear mechanistic reference process instead of a misspecified linear one — which is why it outperforms both. The general statement is therefore that the value of an OT step depends on how well the reference process matches the true dynamics, not on the presence or absence of trajectory inference per se.

The ablation study of Figure 4A–E supports this reading directly. The noloop variant returns the network obtained after a single round of the procedure, without the iterative refinement of Steps 1–3, performing a single round of OT on the burst modes. Its accuracy is accordingly lower than that of the full method — substantially so on BN8, yet it remains above RF on all four benchmark networks (compare Figure 4A–D with Figure 3A).

#### GRN inference for experimental datasets

We applied CardamomOT to the three experimental datasets to obtain GRN structures (Figures S1–S3). The gene sets used for each dataset, and the virtual stimulus nodes included to represent the experimental perturbation applied at *t* = 0 (retinoic acid for Semrau; Dox and Serum for the two successive phases of the Schiebinger protocol), are described in Appendix C.

GRNs inferred across 5 independent runs on each dataset showed mean cosine similarities of 0.777 (Semrau), 0.859 (Kameneva) and 0.798 (Schiebinger) (Figure S9), indicating reproducible overall network structure with residual run-to-run variability. The inferred GRNs are notably sparse (Figures S1A–S3A), with a small number of hub regulators concentrating most of the regulatory activity — a hallmark of biologically realistic network structure [54, 7] that is not always recovered by inference methods without explicit sparsity constraints. Top regulators ranked by outgoing regulatory strength are reported with their main targets in Table S3. The most connected regulators are consistent with those identified in the original studies: Hoxb2, Dnmt3a, Lamb1 and Col4a2 for Semrau (Figure S1C), in agreement with [3], and CHGA among the dominant regulators for Kameneva (Figure S2C), consistent with [38]. Several highly expressed housekeeping genes (RPL30, ATP5F1E) also rank among the top Kameneva regulators (Table S3), which likely reflects their coherent expression across all cells rather than genuine regulatory activity, and illustrates the importance of gene selection discussed in Section 3.

The diagonal coefficients of the inferred GRN matrices, corresponding to auto-regulatory interactions, are systematically non-negligible across all three datasets (Figures S1B–S3B). While often excluded in GRN inference frameworks [8], these terms naturally arise in our setting as a parsimonious explanation for bimodal gene expression dynamics, and are discussed in Section 3. As an internal consistency check, we verified that the inferred targets of the stimulus nodes are broadly consistent with the known biology of each experimental protocol (Appendix C).

These inferred networks serve as the foundation for the perturbation experiments described in Section 2.6.

### 2.4 CardamomOT accurately reconstructs hidden protein vectors, cellular trajectories and velocity fields of simulated data

Unlike existing trajectory inference methods that model dynamics directly on mRNA levels using simplified diffusion processes [15] or linear systems [25], CardamomOT constructs trajectories at the protein level, explicitly accounting for the highly bursty nature of transcription and the slower, more stable dynamics of proteins. We evaluate the benefits of this trajectory reconstruction on three complementary levels:

- the accuracy of the predicted **velocity fields** on cross-sectional data (Figure 5), evaluated in both transcript and protein space;
- the accuracy of the predicted **cell-to-cell couplings** on true trajectory data when simulated cells are not virtually killed at each timepoint (Figure S7);
- the similarity of the hidden protein levels reconstructed by the method compared to the simulated ones, which are not used by the algorithm (Figure S6).

**Figure 5.**
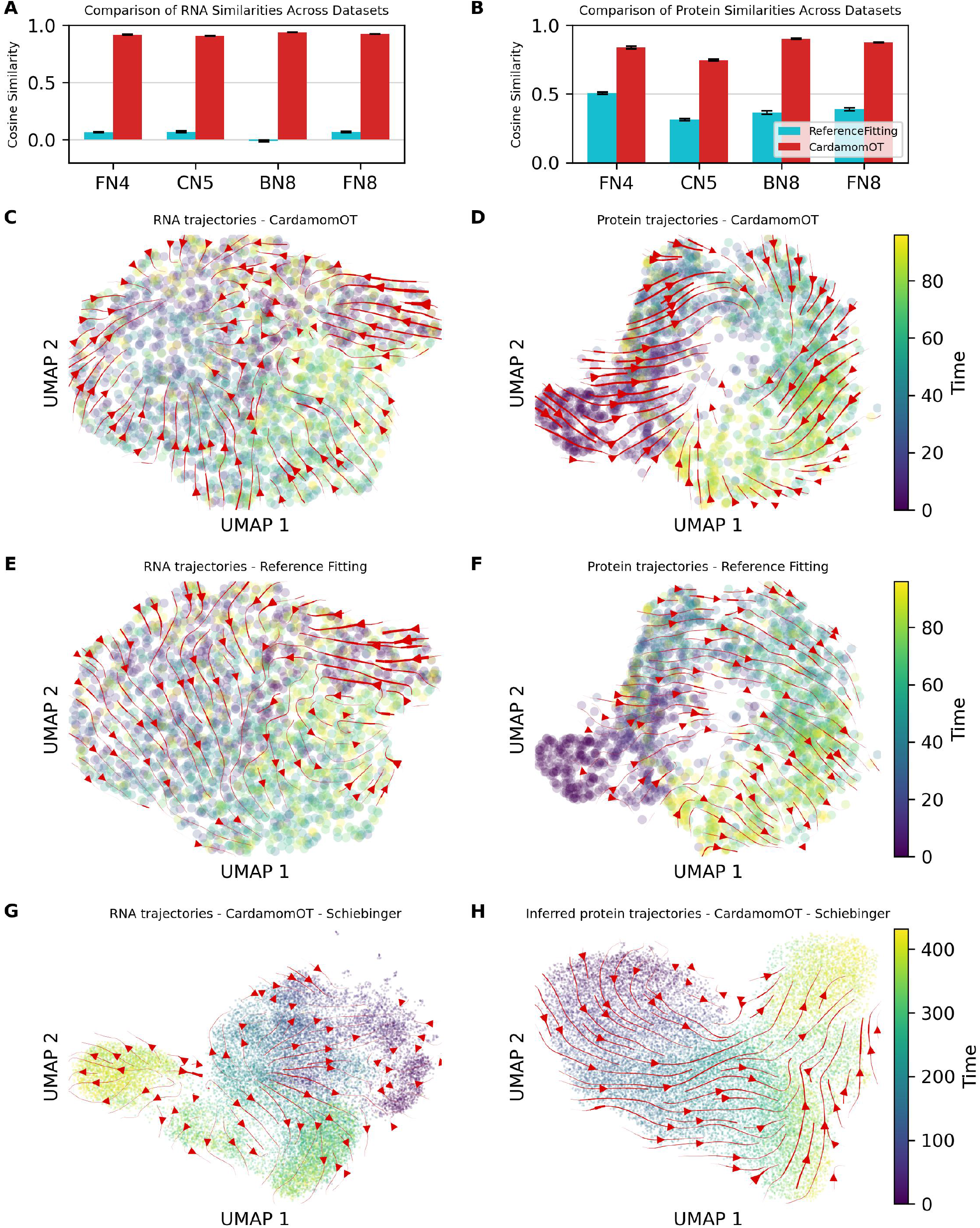
Evaluation of the mRNA and protein velocity fields reconstruction on cross-sectional data for CardamomOT and Reference Fitting. (A–B) Weighted cosine similarity between the velocity fields predicted by CardamomOT and Reference Fitting and the mechanistic ground-truth velocity field, for mRNA (A) and protein (B), on four benchmark networks (BN8, FN8, CN5, FN4). Bars show the mean over 5 independent simulations; error bars indicate the standard error of the mean. (C–F) UMAP embeddings of RNA (C,E) and protein (D,F) expression of CN5 benchmark cells, overlaid with velocity stream plots from CardamomOT (C–D) and Reference Fitting (E–F). (G–H) UMAP of RNA (G) and inferred protein (H) data from the Schiebinger dataset, with CardamomOT velocity stream plots.

#### Velocity fields on cross-sectional data

For the mechanistic model, the expected instantaneous velocity fields for mRNAs and proteins at a given state (*M, P*) are given by the velocity fields *v*_0_ and *v*_1_ described in Eqs. (6) and (5), respectively. For CardamomOT, both velocity fields are evaluated using the inferred parameters (*θ, d*_0_, *d*_1_, *k*_1_) and the reconstructed protein trajectories. For RF [25], the mRNA velocity is given by the instantaneous drift of the calibrated linear Ornstein–Uhlenbeck process, 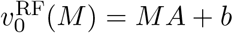, where *A* and *b* are the inferred generator matrix and bias; the protein velocity is derived by applying the same linear operator to the protein values used as a proxy. We evaluate the agreement of each predicted velocity field with the mechanistic reference using a weighted cosine similarity, where weights are given by the product of the norms of the two velocity vectors.

RF’s mRNA-based velocity field shows poor agreement with the mechanistic reference, with cosine similarities close to zero across all benchmark networks (Figure 5A). This reflects an intrinsic limitation: mRNA levels are dominated by stochastic transcriptional bursting, and a global linear model cannot capture the local, protein-dependent regulation that drives gene expression changes. The resulting velocity field is essentially a uniform flow in UMAP space (Figure 5E), bearing little resemblance to the structured, GRN-driven dynamics of the system. In contrast, CardamomOT achieves cosine similarities exceeding 0.8 across all networks (Figure 5A), and the corresponding UMAP stream plots (Figure 5C) show velocity fields with coherent, biologically interpretable structure.

The disparity between methods is also visible at the protein level. Proteins evolve more slowly and deterministically than mRNAs, making their dynamics less sensitive to transcriptional noise and therefore more tractable for trajectory inference. When RF’s linear dynamics are applied to protein space, the agreement with the mechanistic reference clearly improves (cosine similarity ~ 0.25–0.4, Figure 5B), suggesting that even simplified linear models can partially capture protein-level trends. The resulting velocity field is nevertheless still close to a uniform flow in UMAP space, and poorly reflects the true temporal evolution of cells (Figure 5F).

By contrast, CardamomOT— which directly infers protein trajectories and uses the mechanistic model to predict protein velocities — achieves higher cosine similarities (again exceeding 0.75, Figure 5B), with protein UMAP stream plots (Figure 5D) showing well-structured trajectories consistent with the underlying GRN dynamics (in particular a correct orientation of the cycle across timepoint).

#### Coupling accuracy on true trajectory data

To complement this analysis, we also evaluated both methods on simulated datasets where true cell-to-cell correspondences are known, i.e. where individual cells are tracked across consecutive timepoints (Figure S7). In this setting, we can directly assess the quality of the inferred couplings by measuring the expected distance between predicted and observed descendants, with the same metric as in [55]. This metric is computed separately for mRNA (with a log(1 + *x*) transformation to reduce the influence of highly expressed genes) and for proteins (already rescaled in [0, 1]).

CardamomOT achieves consistently lower descendant distances than RF on mRNA (Figure S7A) and proteins (Figure S7B), confirming that the mechanistic coupling provides more accurate cell-to-cell correspondences.

#### CardamomOT reconstructs protein trajectories from transcriptomic snapshots

A key feature of CardamomOT is its ability to infer protein trajectories that are never directly observed in scRNA-seq data. To evaluate this capability on simulated datasets, where ground truth protein values are available, we compared the reconstructed protein trajectories to the true simulated ones across all benchmark networks (Figure S6). The inferred protein UMAP embeddings closely reproduce the structure of the reference trajectories in all cases, capturing both the global topology of the differentiation process and the temporal ordering of cells. This demonstrates that CardamomOT can reliably recover simulated hidden protein dynamics from transcriptomic snapshots alone.

### 2.5 The calibrated model regenerates cellular trajectories consistent with experimental data

A distinguishing feature of CardamomOT is its use as a *generative model* : once calibrated on temporal snapshots, the model (with its inferred GRN, mechanistic parameters and protein dynamics) should be able to simulate new datasets that closely resemble the original data. This generative capability is critical for two reasons: (i) it provides a stringent test of whether the model has captured the essential dynamics of the system, and (ii) it enables *in silico* experimentation through perturbation analysis (Section 2.6).

To validate this capability, we simulated new temporal snapshots starting from *t* = 0 and compared them to the original experimental data using the same metrics as in Section 2.2. Despite the complexity of the inference procedure, the fully calibrated model reproduces the experimental data with a quality comparable to the NB mixture model alone:

- **Low-dimensional embeddings:** Original and simulated cells overlap extensively in joint mRNA UMAP space (Figure 6A–F), confirming that the mechanistic model, driven by the inferred GRN and protein dynamics, captures the global structure of the gene expression landscape while exhibiting realistic stochastic variability. This analysis was repeated using the trajectory-preserving method PHATE [56], providing qualitatively consistent visualization of the global trajectory structure (Figure S10).
- **Marginal distributions and correlations:** The marginal distributions and pairwise gene expression correlations of simulated data are in close agreement with those of the NB mixture model at all timepoints (Figures S4 and S5; *>* 0.95 correlation of gene pair correlations across all datasets), demonstrating that the GRN-driven simulation does not distort the statistical structure captured at the preprocessing stage. The mild underestimation of the strongest correlations observed in Figure S5 is thus inherited from the conditional independence assumption of the NB mixture, and not introduced by the mechanistic model.
- **Regulatory gene dynamics:** Temporal profiles of key regulators show close agreement between original and simulated data in both mean expression and variance (Figure 6J– O). This is a particularly stringent test: errors in GRN inference typically manifest as incorrect dynamics for highly connected genes [32], so this agreement is consistent with the inferred GRN capturing genuine regulatory relationships.
- **Cell type proportions:** Cell type compositions are highly consistent between original and simulated data across all timepoints (Figure 6G–I), confirming that the mechanistic model reproduces the temporal evolution of major cell populations. Nevertheless, for the Kameneva and Schiebinger datasets, some intermediate or rare populations (in particular the “Intermediate” cells in Kameneva and the trophoblasts in Schiebinger) are less well captured, leading to noticeable discrepancies in their inferred proportions. These cell types correspond to subtle variations in expression patterns that are difficult to represent with a statistical model using only two modes per gene, as further discussed in Section 3.

**Figure 6.**
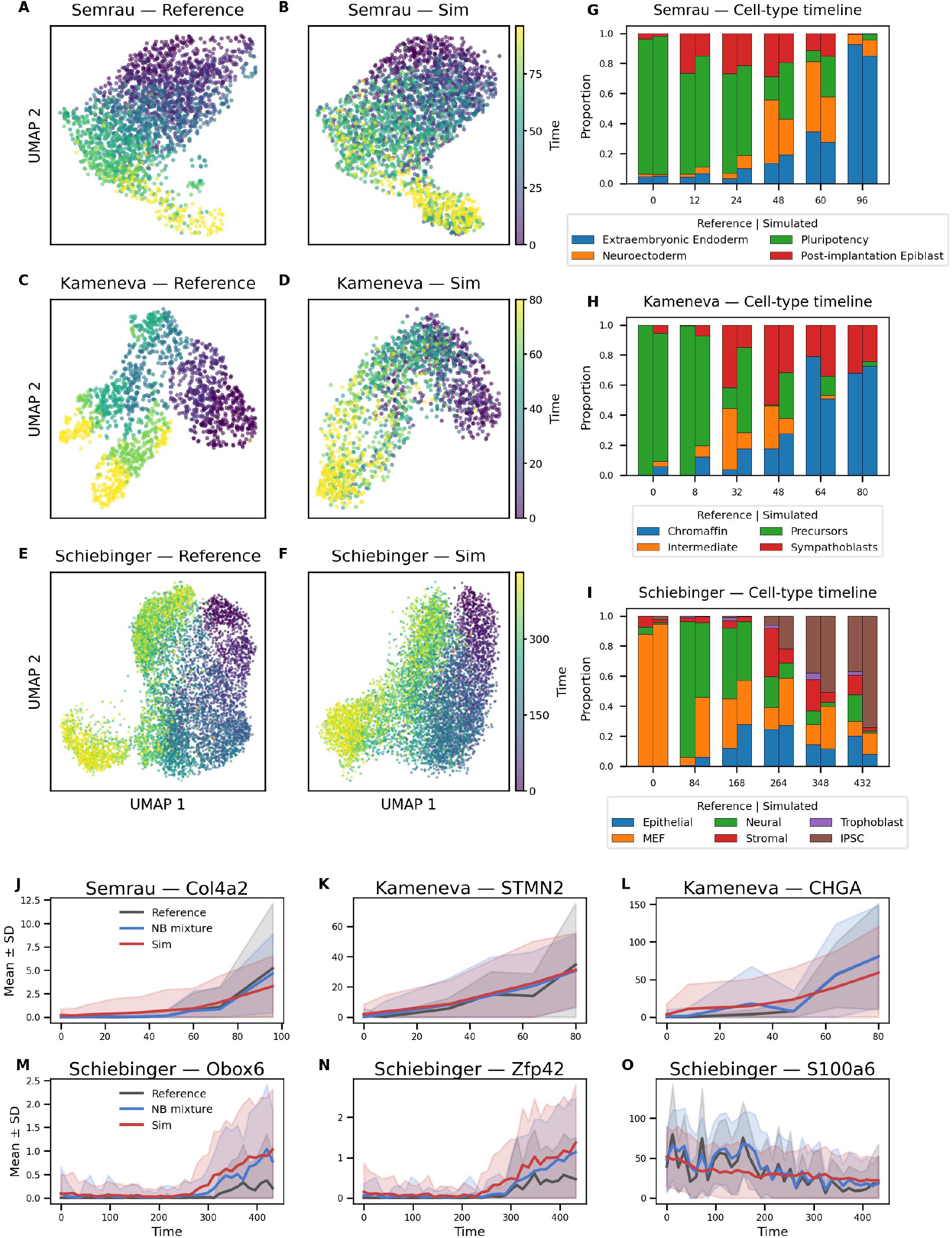
CardamomOT performs well as a generative model of protein and mRNA distributions. UMAPs of reference (A,C,E) and simulated (B,D,F) data for 3 differentiation datasets. Cell type composition for reference and simulated data across timepoints (G–I). Cell type composition was inferred for simulated data by a random forest model trained on reference data. (J–O) Temporal profiles of mean *±* SD gene expression across all cells for selected key regulators, comparing reference data (gray), Negative Binomial reconstruction (blue) and model simulations (red), for Col4a2 in the Semrau dataset (J), STMN2 and CHGA in the Kameneva dataset (K–L), and Obox6, Zfp42 and S100a6 in the Schiebinger dataset (M–O).

These results demonstrate that the mechanistic model inferred by CardamomOT constitutes a generative model of single-cell transcriptomic dynamics and supports its use for predictive perturbation analysis.

### 2.6 In silico perturbations predict some observed experimental outcomes and identify potential key regulators

A central objective of GRN inference is to assess whether the inferred network can predict the effects of unseen genetic perturbations on cell fate decisions. The calibrated CardamomOT model enables such analyses by simulating the system under controlled perturbation scenarios. We considered two types of perturbations: (i) gene knockout (KO), where a gene’s protein level is fixed at zero and its basal parameter *β*_*i*_ is set to −1000, and (ii) gene overexpression (OV), where a gene’s protein level is fixed at one (in the rescaled units of Equation (5)) and its basal parameter *β*_*i*_ is set to +1000 (this value ensuring that it overwhelms the effect of any other regulatory interaction).

For each perturbation, we generated simulated temporal snapshots using the same time points and sampling scheme as in the original datasets. We quantified perturbation effects by measuring changes in cell-type proportions, with cell identities assigned using the Random Forest classifiers described in Section 2.2. To isolate the contribution of the inferred GRN, we compared experimental data to: (i) wild-type simulated data (*sim WT* in Figure 7); (ii) simulated data with the perturbation applied from *t* = 0 (*Sim perturb*); and (iii) wild-type simulated data in which only the expression level of the perturbed gene was replaced by its value under perturbation (*Sim single*).

**Figure 7.**
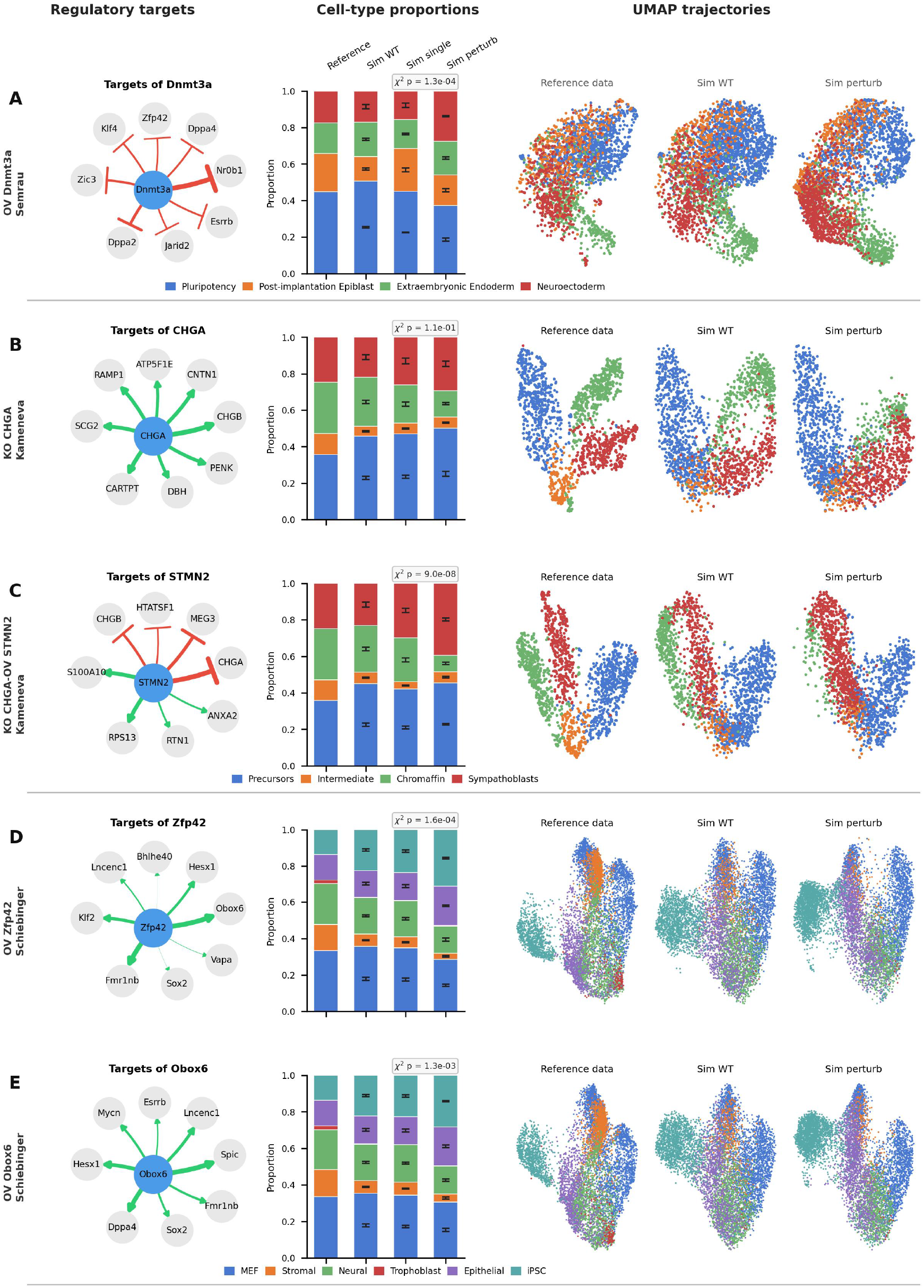
In silico perturbations using the inferred GRNs can recapitulate experimentally observed effects. For each perturbation, three panels are shown: (left) the regulatory network centered on the perturbed gene, showing its top predicted targets with activation (green) and inhibition (red) edges; (middle) cell type proportions comparing reference data, wild-type simulation (Sim WT), simulation with only the perturbed gene modified (Sim single) and perturbed simulation (Sim perturb); (right) UMAP trajectories for reference data, wild-type and perturbed simulations, colored by cell type. The *χ*^2^ test was computed on the base simulated dataset shown here; error bars show the standard deviation of cell-type proportions across *N* = 5 independent stochastic replicates.

Condition (iii) was included to assess whether the observed effects could be explained solely by changes in the expression of the perturbed gene, for example if that gene acts as a strong marker of a given cell type.

To further quantify these differences, we computed *χ*^2^ statistics comparing the cell-type distributions of Sim perturb and Sim single from the base simulated dataset shown in Figure 7. Because the number of simulated cells differs by orders of magnitude across datasets (from a few thousand for Semrau and Kameneva to ~ 250,000 for Schiebinger), using the raw cell counts would make the *χ*^2^ statistic strongly dependent on dataset size, so we rescaled the compared proportions to a fixed, dataset-independent pseudo-count of 100 cells per cell type before computing it. We emphasize that this rescaled statistic is used as a comparative measure of compositional separation rather than as a formal inferential test on the underlying biological system. Variability across independent stochastic replicates is reported separately as error bars in Figure 7. All perturbations except CHGA KO produced compositional shifts larger than the run-to-run variability observed across stochastic replicates, supporting that the predicted effects are not solely driven by marginal changes in the perturbed gene expression but arise from the propagated regulatory interactions encoded in the inferred GRN.

#### ES differentiation (Semrau)

Using the GRN inferred from the ES differentiation time series, we evaluated the effect of Dnmt3a overexpression. The model predicts that Dnmt3a OV substantially alters cell fate distribution, with a marked increase in the proportion of neuroectoderm cells at the expense of pluripotent cells compared to the wild-type simulation (Figure 7A). The predicted regulatory targets of Dnmt3a are predominantly inhibitory, including key pluripotency regulators such as Esrrb, Zfp42 and Klf4, as well as lineage-associated factors such as Hoxb2 and Zic3, consistent with a broad repressive role on the pluripotency transcriptional program.

This result is particularly interesting from a mechanistic perspective. Unlike TFs that directly contact the transcription machinery, Dnmt3a is a de novo DNA methyltransferase that acts indirectly on gene regulation through CpG methylation — a well-documented epigenetic mark generally associated with gene expression silencing [57]. The predominantly inhibitory interactions inferred by CardamomOT for Dnmt3a are consistent with its known epigenetic function, illustrating that the GRN model is capable of capturing indirect regulatory effects mediated by epigenetic mechanisms and is not restricted to direct physical interactions between TFs and their targets.

#### Sympathoadrenal differentiation (Kameneva)

The analysis in [38] identified four important regulators: CHGA, STMN2, CHGB and HMGB2. We quantified the effect on the proportion of chromaffin cells and sympathoblasts, the principal cell types in sympathoadrenal development, associated with the perturbation of each of these genes alone or in combination. Among the tested perturbations, KO of CHGA alone produced a moderate reduction in the proportion of sympathoblasts, of an amplitude comparable to the run-to-run variability of the simulations (Figure 7B). The combined perturbation consisting of CHGA KO and STMN2 OV, by contrast, markedly amplified this effect, nearly abolishing the emergence of sympathoblasts in the simulations (Figure 7C). This contrast between the single and combined perturbations indicates that the inferred GRN captures non-trivial combinatorial effects on lineage commitment, rather than a simple additive contribution of each perturbed gene. We emphasize that, unlike the Obox6 and Zfp42 predictions discussed below, these two perturbations have not been tested experimentally and should be read as model-generated hypotheses.

#### MEF-to-iPSC reprogramming (Schiebinger)

In [15], the authors reported that OV of Obox6 and Zfp42, two TFs not typically included in standard reprogramming cocktails, enhances the efficiency of MEF-to-iPSC conversion. To assess whether CardamomOT recovers this behavior, we inferred a GRN from the reprogramming time series and simulated the effect of Obox6 and Zfp42 OV.

Both perturbations consistently increase the proportion of iPSC-like cells in the simulations compared to control conditions (Figure 7D,E), in line with the experimental observations of [15], with a similar magnitude of effect (for Obox6 OV we measure an increase from ~ 20% to ~ 40% of iPSC-like cells). Importantly, this effect emerges solely from the temporal scRNA-seq data used for inference, without incorporating any prior knowledge about the function of these factors.

The predicted regulatory targets provide a mechanistic interpretation of this behavior. Zfp42 OV activates several pluripotency-associated genes, including Esrrb and Dppa4 (Figure 7D), both well-known markers of the naive pluripotent state. Obox6 OV similarly activates Gata3, Lncenc1 and Mycn, and notably upregulates Zfp42 itself (Figure 7E), suggesting a positive feed-back loop between the two factors that may reinforce the iPSC-like state. Rather than inducing broad lineage rewiring, both perturbations thus appear to strengthen specific components of the pluripotency transcriptional program already present in the data.

Taken together, these perturbation prediction analyses support the use of CardamomOT as a framework for generating quantitative, mechanistically interpretable perturbation hypotheses across distinct biological systems. While experimental validation remains essential, the consistency observed across multiple datasets supports the relevance of the inferred networks for studying cell fate control. Interestingly, the four regulators perturbed in silico above (Dnmt3a, CHGA, Obox6, Zfp42) represent only a fraction of the candidate hub regulators inferred across datasets (Table S3), which also includes regulators with little or no established role in the corresponding process — e.g. the lncRNA HAND2-AS1 (Kameneva) or the poorly annotated Dmrtc2 and Gm26917 (Schiebinger). Their co-occurrence with validated regulators in the same ranked output illustrates the potential of CardamomOT to generate testable hypotheses about uncharacterized genes, though such candidates should be weighed against biological plausibility — a subset of top Kameneva hits (RPL30, ATP5F1E) instead reflect housekeeping expression rather than genuine regulatory activity (Section 2.3).

## 3. Discussion

Taken together, our results demonstrate that CardamomOT provides a complete pipeline for mechanistic analysis of temporal scRNA-seq data. Starting from raw count matrices and timepoint labels, the method (i) infers mechanistic parameters and identifies discrete regulatory states (Section 2.2), (ii) reconstructs both GRN structure and hidden protein trajectories through an iterative refinement procedure (Sections 2.3 and 2.4), (iii) generates a calibrated model that can simulate realistic cellular dynamics (Section 2.5), and (iv) predicts the effects of unseen genetic perturbations (Section 2.6).

### A unified framework for GRN inference and trajectory reconstruction

Most existing approaches treat GRN inference and trajectory inference as separate problems: GRN inference methods typically assume static or quasi-static cellular states, while trajectory inference methods focus on reconstructing cell-to-cell transitions without explicitly modeling gene regulation. CardamomOT unifies these perspectives through an iterative procedure that alternates between reconstructing protein trajectories given a GRN (Step 1) and updating the GRN structure given the trajectories (Step 2), until convergence.

This iterative refinement is essential for achieving high inference accuracy: removing the iterative loop substantially degrades performance (Figure 4), confirming that the GRN and trajectories contain complementary information that must be jointly optimized. At the same time, the method converges to similar GRN structures regardless of initialization in our benchmarks (Figure 4A–E), suggesting that the temporal structure of the data provides sufficient constraints to escape local optima for the network topologies and temporal resolutions tested here.

The mechanistic foundation of CardamomOT also enables trajectory reconstruction at the protein level rather than the mRNA level. As shown in Section 2.4, mRNA-based velocity fields exhibit poor agreement with the underlying mechanistic dynamics due to the highly stochastic, bursty nature of transcription, whereas protein dynamics are slower and more deterministic, providing a more stable substrate for trajectory inference. By explicitly modeling the conditional distribution of mRNA given protein levels, CardamomOT captures both bursty transcriptional dynamics and relatively smooth, GRN-driven protein trajectories, achieving high cosine similarities with respect to the ground-truth velocity fields (Figure 5B).

### Mechanistic modeling improves interpretability and generalizability

Unlike black-box or purely data-driven approaches, CardamomOT is grounded in an explicit mechanistic model of gene expression derived from biophysical principles. This provides two key advantages. First, the inferred parameters—burst frequencies, burst sizes, and protein degradation rates—are directly interpretable in terms of molecular processes, enabling biological insights beyond network topology alone. Second, by incorporating prior knowledge about kinetic rates and known regulatory interactions, CardamomOT can leverage existing biological information to improve inference, particularly in data-limited regimes.

Robustness to uncertainty in kinetic parameters is a practical consideration for experimental applications. As shown in Figure 4F, CardamomOT maintains high inference accuracy even when protein degradation rates are perturbed by up to *±*50% from their true values, and clearly outperforms baseline methods even with perturbations as large as *±*200–400%. This robustness arises because the iterative procedure jointly optimizes the GRN structure and the inferred trajectories, allowing the method to partially compensate for kinetic misspecification through adjustments in the network topology.

The same mechanistic framework allows CardamomOT to function as a generative model. Once calibrated on experimental data, the model can simulate new datasets that closely reproduce the original observations in terms of marginal distributions, low-dimensional embeddings, and cell-type proportions (Figure 6). This generative capability simultaneously provides a stringent test of whether the model has captured the essential dynamics of the system and enables *in silico* experimentation, including perturbation analysis and prediction of unseen cell fates.

### Perturbation predictions and biological relevance of inferred GRNs

A central objective of GRN inference is to predict how unseen genetic perturbations will alter cellular behavior. CardamomOT addresses this challenge by simulating perturbations directly within the calibrated mechanistic model. In the experimental systems analyzed here, the inferred GRNs are consistent with the perturbation effects for which independent experimental evidence is available, such as the impact of TF overexpression on reprogramming efficiency, and suggest additional regulatory relationships that could be tested experimentally. These results indicate that CardamomOT infers network topology together with regulatory relationships that appear functionally interpretable, although the available validation remains limited to a small number of cases. It should also be noted that inferred GRNs are directly constrained by the information contained in the observed trajectories, which depends both on the nature of the data and on the gene set selected; we return to this point below.

### Optimal transport as a mechanistic framework for trajectory inference

The use of OT for trajectory inference has become widespread in recent years, motivated by the idea that cells follow trajectories that minimize a cost in gene expression space. However, most OT-based methods implicitly assume that cells follow Brownian motion with a constant diffusion rate, corresponding to a Euclidean or squared-Euclidean cost. This assumption lacks biological justification and ignores the mechanistic constraints imposed by gene regulation [23].

CardamomOT extends this framework by introducing a mechanistic cost function that explicitly incorporates the GRN structure. The cost of transitioning between two cells is determined by how well the GRN-predicted burst rates match the observed burst modes, naturally accounting for the fact that gene expression dynamics are driven by protein-mediated regulation rather than passive diffusion. Our approach can be viewed as a generalization of RF to the case where the reference process is a nonlinear mechanistic model instead of a linear Ornstein–Uhlenbeck process, making it suitable for modeling complex regulatory dynamics.

From a theoretical perspective, there is a direct connection between our approach and Wasserstein Lagrangian Flows, which incorporate potential energy constraints into the OT Lagrangian. CardamomOT implicitly defines a Lagrangian derived from the mechanistic model via stochastic calculus (see Appendix B), but avoids the computational burden of parametrizing all characteristics of the process and time-varying measures with neural networks. By discretizing the problem into a sequence of pairwise OT problems constrained by the mechanistic model, CardamomOT achieves computational efficiency while retaining mechanistic interpretability.

### Scalability and identifiability

In our experiments, the computational cost of CardamomOT was not the primary bottleneck: inference on networks of 100 genes and 1000 cells completed in under a minute (Table S2), while inference of approximately 100 genes and tens of thousands of cells completed in about one hour. The true limiting factor is the identifiability of the network parameters. With *n* genes and a fully connected GRN, the number of parameters scales as *n*^2^, *i*.*e* 10^4^ for *n* = 100; beyond this scale, even though running CardamomOT is temporally achievable, the information contained in temporal snapshots may be insufficient to robustly constrain the network structure.

Importantly, from a theoretical perspective, the convergence of our iterative procedure to a global optimum is not guaranteed, as the objective (20) is non-convex in general; the empirical convergence observed throughout our experiments should therefore be understood as convergence to a stationary point, whose quality depends on the temporal structure and richness of the data.

The stochasticity inherent in our iterative procedure nevertheless provides a natural framework to assess identifiability of this optimum. Indeed, the OT coupling sampled at each iteration introduces randomness analogous to the temperature parameter in simulated annealing, so that different initializations may in principle converge to different local optima. In practice, our benchmark experiments (Figure 4A–E) show that the method converges to similar GRN structures regardless of initialization, suggesting that the temporal data provide sufficient constraints for the simulated networks tested here. Inferred GRNs on experimental data show mean cosine similarities of independent runs ranging from 0.777 to 0.859 depending on the dataset, i.e. reproducible overall network structure with residual run-to-run variability (Figure S9). Since these more complex settings do not provide the same constraints, we recommend running inference on 3–5 random seeds for networks *>* 20 genes; high variability in edge weights across runs would signal poor identifiability for that edge. Systematically exploring the variability of optimal networks across different initial couplings would therefore be a principled way to quantify identifiability as a function of network size and data richness, and represents a promising direction for future work.

Addressing identifiability for larger networks will require incorporating external biological constraints. Prior regulatory information—such as TF binding databases, ChIP-seq data, or resources like OmniPath [58] can substantially reduce the effective parameter space. A prior network construction option using the Neko tool [59] via OmniPath is already available in the CardamomOT repository.

A complementary strategy concerns the structure of the basin space. While the number of theoretically possible basins for *n* genes is 2^*n*^, biological systems occupy only a small fraction of this space: not all combinations of per-gene expression modes are accessible, as cell types correspond to coherent regulatory states. It is therefore both possible and desirable to introduce an intermediate step that restricts the basin space using prior biological information—such as known cell types or unsupervised clustering—combined with mixture-of-experts approaches to construct burst-mode vectors from per-gene univariate NB fits. Reducing the effective basin space in this way would simultaneously improve identifiability and computational efficiency.

### Gene selection and its influence on inferred networks

All experimental applications presented here operate on gene subsets rather than the full transcriptome (Appendix C), which is not a neutral choice. Regulatory activity can only be attributed to included genes, so that the influence of an omitted regulator is redistributed among retained genes and inferred edges may summarise indirect regulation through unobserved intermediates. Selecting genes from known markers or previously characterised modules, as for the Semrau and Kameneva datasets, additionally biases the network toward well-characterised regulatory axes: the agreement between inferred hub regulators and those of the original studies (Section 2.3) is thus a consistency check on a pre-selected gene set rather than an unbiased recovery. More fundamentally, the selected gene set should be recognised as a form of prior information in its own right, on the same footing as the prior constraints on individual edges discussed above: it determines which regulatory relationships the method is allowed to express at all — and, when made on statistical grounds alone, which spurious ones it is likely to produce, as illustrated by the housekeeping genes ranking among the top inferred Kameneva regulators (Section 2.3). We therefore recommend making this choice explicit and, where possible, splitting it in two. Prior biological knowledge of the system under study, or the specific hypotheses motivating the analysis, can be used to designate a core set of nodes of interest; the regulatory context of these nodes can then be assembled semi-automatically, by combining curated interaction databases such as those discussed above with data-driven discovery — for instance by using a genome-wide scalable inference method to obtain a coarse-grained approximation of the global network, such as OTVelo [60], and retaining the genes that this approximation places in the neighbourhood of the core set. Automating this construction is the object of ongoing work in our team. This also clarifies the practical form taken by the identifiability ceiling discussed above, which is best understood as a trade-off between the size of the network considered and the amount of prior structure imposed on its edges. The fewer prior constraints one is willing to inject on the structure of *θ*, the smaller the gene set must be — we recommend not exceeding of the order of 100 genes in the fully unconstrained setting; conversely, the more prior structure one accepts, whether through a prior network *θ*^0^ or through zeroed entries, the larger the gene set can be for a comparable level of reliability.

### Diagonal coefficients and auto-regulatory interactions

A specificity of CardamomOT concerns the diagonal coefficients of the inferred GRN matrix, i.e. auto-regulatory interactions. As visible in the inferred networks on experimental datasets (Figures S1–S3), CardamomOT tends to infer non-negligible auto-activations for many variable genes. This is not a numerical artifact: in the mechanistic model, the combination of a positive auto-activation with a negative basal parameter *β*_*i*_ is the most parsimonious way to produce bistability for an isolated gene, and therefore to reproduce transitions between two expression modes. It is thus natural that for any gene exhibiting bimodal dynamics, the model partially attributes this variability to self-reinforcing regulation [34].

Importantly, this tendency is not purely a mathematical convenience. Positive auto-regulatory feedbacks are well documented as a mechanism for stabilizing cell states and enhancing the robustness of cell fate decisions [61]. In this sense, the auto-activations inferred by CardamomOT may reflect genuine biological regulatory structure rather than overfitting. We note that most GRN inference methods explicitly exclude diagonal coefficients from inference [7], which circumvents this question entirely at the cost of ignoring a potentially relevant class of interactions.

In CardamomOT, we deliberately choose not to apply a specific penalty on diagonal coefficients, allowing the model to use auto-activation as an explanation for observed bimodality when this is consistent with the data. In practice, however, we recommend that users of CardamomOT specify in advance whether auto-regulatory interactions are biologically plausible for their system of interest—for instance by setting the corresponding entries of the prior network *θ*^0^ to zero—in order to prevent the model from systematically routing unexplained variability through this pathway.

### Limitations and future directions

Despite its strengths, the current version of CardamomOT has limitations that open avenues for future development.

#### Model assumptions

CardamomOT relies on several simplifying assumptions, including the two-state model of transcription, the bursty regime approximation (*k*_*on*_ ≪ *k*_*off*_), and the use of sigmoids to model GRN-dependent burst rates. While these assumptions are well supported by experimental observations [62, 63], they may not hold universally across all genes and cell types. Notably, an extension to softmax activation functions — which generalize the sigmoid to genes exhibiting more than two expression modes — is already implemented in the CardamomOT package. In practice, this extension introduces a preliminary model selection step in which the optimal number of modes per gene is determined from a user-specified maximum using an AIC criterion; the full inference pipeline is then adapted to handle gene-specific numbers of modes. This comes at the cost of reduced interpretability, since each mode transition is governed by a distinct regulatory network. Further extensions could incorporate even more flexible regulatory functions, such as Hill functions [64, 33] or more complex neural network-based parametrizations [65], albeit with further reductions in interpretability and identifiability.

#### Temporal resolution

CardamomOT requires temporal snapshots at multiple time points, and its performance depends on the temporal resolution. The OT assumption—that between two consecutive timepoints, mode transitions are direct with no unobserved intermediate states—is well justified when timepoints are sufficiently close relative to regulatory timescales. For datasets with sparse or irregular sampling, rapid transient dynamics involving multiple mode transitions may be missed. Extending the framework to accommodate intermediate unobserved switches would improve robustness, albeit at an additional computational cost.

#### Proliferation and death

The current implementation of CardamomOT assumes that cells neither proliferate nor die between consecutive timepoints, so that the marginal constraints of the OT problem are strictly enforced. This assumption is reasonable for many differentiation datasets, but may break down in systems with significant proliferation or apoptosis. Incorporating growth and death rates into our framework is however conceptually straightforward within the OT formalism. When proliferation and death rates can be estimated from external data — for instance from cell cycle markers or DNA content, as in [15] — they can be directly folded into the marginal constraints of the OT problem following the same strategy. When lineage tracing data are available alongside transcriptomic snapshots, the observed lineage tree can be used to deconvolve the contribution of proliferation from that of regulatory dynamics, following an approach analogous to [66]. In the absence of such prior information, unbalanced OT [67] provides a natural relaxation of the marginal constraints, allowing the inferred coupling to account for unequal cell numbers across timepoints without requiring explicit knowledge of proliferation or death rates. These extensions lie beyond the scope of the present work, but are actively being integrated into the CardamomOT package.

#### Multi-omic extensions

Beyond the transport formalism itself, the mechanistic model underlying CardamomOT can also be enriched to accommodate richer data modalities. In its current form, it describes gene expression dynamics at the level of mRNA and protein, and is thus naturally suited to transcriptomic data, possibly augmented by proteomic measurements. Multi-omic single-cell technologies are now increasingly available, jointly profiling transcription and chromatin accessibility (e.g. scATAC-seq), DNA methylation, or histone modifications. These data provide complementary mechanistic information: chromatin accessibility reflects the regulatory state of promoters and enhancers, and is directly related to the promoter-switching dynamics modeled here; methylation provides information about longer-term epigenetic memory. Extending our mechanistic framework to jointly model transcription, chromatin dynamics, and other epigenomic layers is a major direction for future work. Our inference approach — grounded in the Schrödinger problem and relying on state-of-the-art OT tools — is well suited to be extended to such richer mechanistic models, and we anticipate that this line of development will substantially expand the scope and biological resolution of mechanistic single-cell inference.

In summary, CardamomOT provides a principled, mechanistically grounded framework for joint GRN inference, trajectory reconstruction, and generative modeling from temporal snapshots of single-cell transcriptomic data. By explicitly modeling the stochastic dynamics of gene expression and integrating OT theory with mechanistic modeling, it advances our ability to decode the regulatory programs that govern cellular differentiation and reprogramming. Beyond inference, the calibrated model functions as an in-silico representation of the biological system: it can simulate realistic single-cell datasets, predict the effects of unseen genetic perturbations, and generate mechanistically interpretable hypotheses about cell fate control — capabilities that are not jointly addressed by the comparator methods considered here.

We anticipate that CardamomOT will be most powerful in biological systems where prior knowledge is available to constrain the inference: systems with characterized protein kinetics, known regulatory modules, or existing network resources such as OmniPath [58]. As single-cell atlases of well-studied differentiation and reprogramming systems mature, such settings are increasingly common, and CardamomOT could become a useful reference tool for mechanistic single-cell analysis, bridging the gap between data-driven trajectory inference and biophysically grounded models of gene regulation.

## A. Detailed interpolation procedure

We describe how the candidate protein vector 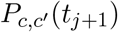 is computed for each pair of cells (*c, c*′) at consecutive timepoints (*t*_*j*_, *t*_*j*+1_). The goal is to integrate the protein ODE (5) between *t*_*j*_ and *t*_*j*+1_, while allowing for a single switch of the driving burst mode.

### Switching time

For each source cell *c* and each gene *i*, we introduce a *switching parameter α*_*c,i*_ ∈ (0, 1) that encodes the fraction of the interval [*t*_*j*_, *t*_*j*+1_] elapsed before the mode of gene *i* transitions from the basin of *c* to that of *c*′. Formally, the protein of gene *i* evolves under basin *z*_*c*_ on [*t*_*j*_, *t*_*j*_ + *α*_*c,i*_Δ*t*] and under basin 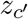 on [*t*_*j*_ + *α*_*c,i*_Δ*t, t*_*j*+1_], where Δ*t* = *t*_*j*+1_ − *t*_*j*_.

### Explicit interpolation formula

Since (5) is a linear ODE with piecewise constant driving term, it can be integrated in closed form on each sub-interval. For gene *i*, denoting by 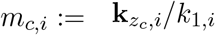 and 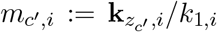 the normalized basin modes at *t*_*j*_ and *t*_*j*+1_, respectively, we obtain

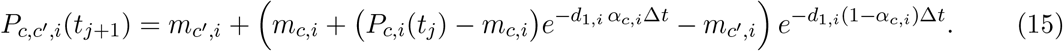

### Use of observed mRNAs as mode proxies

To better reflect the stochasticity of mRNA in the full model (2), we replace the normalized basin modes *m*_*c,i*_ by observed (normalized) mRNA counts 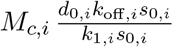 as proxies for the true modes. This provides a richer, cell-specific signal while introducing only a mild approximation (gene expression noise and mode uncertainty are partially conflated). In practice, this proxy yields more realistic protein trajectories, especially when cells within the same basin show broad mRNA distributions.

### Iterative update of switching parameters

The switching parameters *α*_*c,i*_ are updated at each iteration of the main loop. For a given cell *c* at time *t*_*j*_:

1. For each gene *i*, we compute 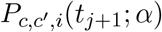 using (15) at a grid of candidate values α ∈ *{*1*/n*_pas_, 2*/n*_pas_, …, 1*}*.
2. We evaluate 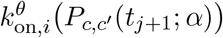 for each candidate *α* under the current GRN *θ*.
3. We set *α*_*c,i*_ to the smallest *α* such that the predicted burst rate is closer to the target mode 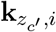 than to the source mode 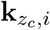, i.e. the earliest time at which the trajectory commits to the new basin.

Genes for which the difference between source and target burst rates is very small (below a threshold *δ* := 0.1) are left unchanged, as their switching time cannot be reliably identified from the data. This update scheme ensures that *α*_*c,i*_ progressively encodes the effective transition timing as the GRN estimate improves.

### Interpretation of the interpolation

The interpolation described above can be interpreted as an approximate bridge for the mechanistic model (2): by conditioning on observed mRNA states at both endpoints and integrating the protein equation with a single mode switch at time *α*Δ*t*, we obtain a deterministic approximation of the most probable protein path between the two observations. The iterative update of *α* refines this approximation alongside the GRN.

### Regime of validity

Eq. (15) is not a numerical discretization of a nonlinear ODE. Within [*t*_*j*_, *t*_*j*+1_] we replace 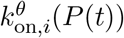 by a piecewise-constant driving term taking the two endpoint basin modes with a single switch at *t*_*j*_ + α_*c,i*_Δ*t*; Eq. (5) is then linear and Eq. (15) is its exact closed-form solution. The approximation therefore lies in the substitution — the regulatory feedback is frozen within the interval — and in the assumption that a single switch per gene suffices to describe it, together with the mRNA-proxy approximation stated above; the grid over *α* is a search over switching times, not a time-stepping scheme.

This piecewise-constant treatment concerns only the construction of candidate protein paths between two observed timepoints. The nonlinear, GRN-driven structure is retained wherever the network is estimated or used: in the OT cost of Eq. (8), the GRN regression of Eq. (10) and the label refinement of Eq. (11); in the kinetic recalibration of Section 1.2.3, where the full field *v*_1_ is integrated with an adaptive-step solver; and in the generative simulations, which sample the complete PDMP (2). The linearity of Eq. (15) is thus a local device for generating transport candidates, not a linearization of the inferred dynamics.

Both approximations hold in the metastable, bursty and adiabatic regime already required by the model of Section 1.1: within a basin the driving mode is close to constant, and inter-basin transitions are rare and fast. Outside this regime the discrete-basin representation itself — the mixture (4) and its labels *z* — ceases to describe the data, since cells can no longer be assigned to a small number of promoter-activity states. The interpolation therefore adds no restriction beyond those the mechanistic model already requires: the two approximations share a regime of validity and break down together. The remaining case, several switches within one interval, is the sampling-resolution limitation discussed in Section 3.

### Practical choice of *n*_pas_

The number of grid points *n*_pas_ used to search for the switching time *α*_*c,i*_ (default 25, and in practice further increased automatically so that at least one grid point falls within each unit of the inter-timepoint interval Δ*t* = *t*_*j*+1_ − *t*_*j*_) has little practical influence on the inferred trajectories once it is not chosen pathologically small. Because Eq. (15) varies smoothly in *α*, the interpolation error introduced by a coarser grid is itself smooth. Moreover, *α*_*c,i*_ is re-estimated at every iteration of the main EM loop (Section 1.2.2) jointly with the GRN, so any residual discretization error from an early, coarse estimate is corrected over subsequent iterations. A sensitivity analysis under refinement of *n*_pas_ confirms this (Figure S11): AUPR plateaus from *n*_pas_ ≈ 4 onwards on all four benchmark networks, and only the extreme value *n*_pas_ = 2 — which removes essentially all timing resolution within an interval — degrades accuracy.

## B Link with other model learning approaches

We briefly explain how CardamomOT fits into the broader family of Schrödinger- and OT-based trajectory inference methods, and how it relates to RF [25] and Wasserstein Lagrangian Flows [24].

### B.1 The Schrödinger problem as a template for trajectory inference

Let *Q* be a reference path measure on gene expression trajectories over [*t*_1_, *t*_*N*_]. Given empirical marginals 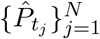 from a time-stamped scRNA-seq dataset, the (multimarginal) Schrödinger problem seeks the path measure closest to *Q* that matches the observed marginals:

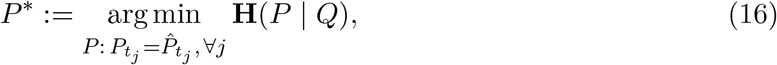

where **H**(*P* | *Q*) is the relative entropy (KL divergence) of *P* with respect to *Q*. When *Q* = *W*^*σ*^ is a Brownian motion with diffusion *σ*, (16) reduces to the entropic OT problem used in [15]: its solution *P* ^*^ describes the most likely cellular trajectories under a free-diffusion prior. The two-marginal version between consecutive timepoints yields the classical Sinkhorn algorithm.

In the diffusion setting, (16) can be reformulated as a variational problem with Lagrangian

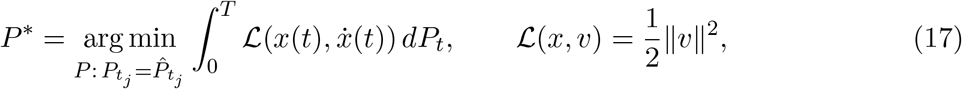

which is the dynamic Benamou–Brénier formulation of OT [22]. The choice of reference process *Q* thus encodes a prior on cellular dynamics.

### B.2 Reference Fitting and Wasserstein Lagrangian Flows

#### Reference Fitting

RF [25] replaces the Brownian reference by a linear Ornstein–Uhlenbeck (OU) process *U*^*θ*^, whose drift *A*^*θ*^*x* is parametrized by a network matrix *θ*. The corresponding problem reads

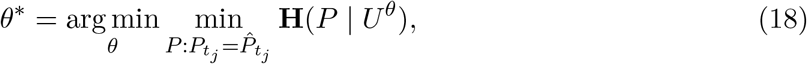

where the inner minimization is again a Schrödinger problem (solved pairwise with Gaussian Sinkhorn), and the outer minimization fits the linear drift. This restricts the dynamics to linear systems and does not capture nonlinear GRN behaviour.

#### Wasserstein Lagrangian Flows

WLF [24] instead modify the Lagrangian by adding a potential energy term *V* (*x*):

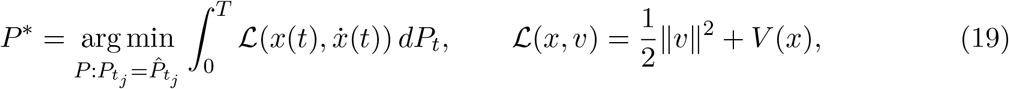

where *V* acts as a prior on the energy landscape of cells. All components (time-varying measures and drift) are learned jointly through neural networks, providing a flexible but computationally demanding framework.

### B.3 CardamomOT as a mechanistic Schrödinger problem

In CardamomOT, the reference process is the GRN-driven model (2), which belongs to the general class of PDMPs. This nonlinear, non-diffusive process induces a Markovian path measure on the basin space via the phenomenological model introduced in Section 1.1: the basin label *z*_*c*_(*t*) follows a Markov chain whose transition rates are determined by *θ* through a complex relationship involving the computation of mean first passage times between basins for protein dynamics, as explored in detail in previous work [44]. We denote this path measure by *Q*^*θ*^.

The Schrödinger problem on the basin space thus becomes

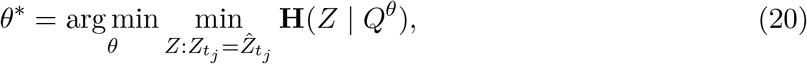

Where 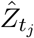 is the empirical distribution of basin labels at time *t*_*j*_, obtained in the preprocessing step. This is structurally analogous to (18), with the linear reference *U*^*θ*^ replaced by the mechanistic Markov chain *Q*^*θ*^ on the space of basins, whose rates depend on the non-linear underlying dynamics on proteins.

### B.4 The EM loop as an iterative minimizer of (20)

The alternating optimization in CardamomOT (Steps 1–3 of Section 1.2.2) can be viewed as a coordinate-descent algorithm for (20).

Minimizing over *Z* with fixed *θ* (Step 1) corresponds to computing a Schrödinger bridge with a structured reference process *Q*^*θ*^ [68], which provides a mechanistic justification for the cost in (8). Indeed, basin dynamics can be viewed as the evolution of the effective activation state 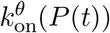 and are therefore driven by the underlying protein dynamics. In the metastable regime, where proteins follow the deterministic limit within each basin and undergo rare stochastic transitions between basins, this induces an effective dynamics on the space of basins whose associated action yields the cost in (8).

Minimizing over *θ* with fixed *Z* (Step 2) is a regression step that fits the GRN to basin transitions, equivalent to maximizing a path likelihood for *Q*^*θ*^ along sampled trajectories. Step 3 updates the empirical marginals 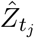 by refining basin labels consistently with the current *θ*.

Together, these steps define an iterative scheme converging toward a stationary point of (20), analogous to Iterative Proportional Fitting (Sinkhorn) but with a mechanistic reference process on the latent basin space.

### B.5 Summary

**Table 1.** Positioning of CardamomOT within the Schrödinger/variational OT family. The key distinction is the use of a mechanistic, non-diffusive reference process defined on the latent basin space rather than on mRNA space.

| Method | Reference process $Q$ | Lagrangian modification | Ambient space |
| --- | --- | --- | --- |
| WOT / Sinkhorn [15] | Brownian motion | None | mRNA |
| RF [25] | Linear OU | None | mRNA |
| WLF [24] | Brownian motion | $V(x)$ | mRNA |
| **CardamomOT** | **Non-linear PDMP (2)** | **None** | **basin** |

In summary, CardamomOT can be seen as a mechanistic generalization of RF, using a nonlinear GRN-driven PDMP as reference instead of a linear SDE. The connection to WLF arises through the Lagrangian interpretation: the mechanistic GRN effectively defines an energy landscape that constrains inferred trajectories, but here this landscape is derived from biophysical modeling rather than learned from data.

## C Details on GRN inference for experimental datasets

This appendix provides additional detail on the gene selection procedure, the stimulus nodes introduced in each experimental dataset, and the consistency between their inferred targets and the known biology of each protocol. It complements the summary given in Section 2.3.

### Gene selection

For the Semrau and Kameneva datasets, all available cells were used directly on the subset of genes identified in [32] and [38], respectively. For the Schiebinger dataset, genes were selected as a combination of markers identified in the original study [15] and dynamically variable genes across timepoints, the latter being identified using a weighted measure combining per-gene entropy and Wasserstein distances between consecutive timepoint distributions, following [38].

### Stimulus nodes

Following the simulation framework described in Section 2.1, a virtual stimulus gene is included in each inferred network to represent the experimental perturbation applied at *t* = 0. For the Semrau dataset, this corresponds to the addition of retinoic acid, which triggers ES cell differentiation [3]; for the Schiebinger dataset, it represents the doxycycline-induced expression of the OSKM reprogramming factors [15]. For the Kameneva dataset, where cells are ordered by pseudotime rather than experimental time and no clear exogenous stimulus is applied, a stimulus gene is nonetheless included to push cells out of their initial equilibrium state, but its contribution to the network is strongly penalized via the prior regularization so that it does not dominate the inferred regulatory interactions.

Because the Schiebinger protocol involves two successive phases — a serum/LIF medium in addition to an initial doxycycline-induced expression of the OSKM reprogramming factors (days 0– 8), followed by withdrawal of doxycycline (days 8–18) [15] — we introduced two distinct stimulus nodes rather than a single averaged one: a Dox node active during the induction phase (days 0–8), and a Serum node active throughout the experiment (days 0–18).

### Consistency of inferred stimulus targets with known biology

The stimulus node inferred for the Semrau dataset corresponds to retinoic acid (RA) and predominantly represses naive pluripotency regulators such as Esrrb, Klf2 and Klf4, while activating Dnmt3a and Jarid2 (Figure S1F), broadly consistent with the known pro-differentiation effect of retinoic acid on mouse embryonic stem cells. For the Schiebinger dataset, the inferred targets of Dox include the lack of activation of the late pluripotency marker Dppa5a and the epithelial gene Cldn7, consistent with the early phase of reprogramming preceding the mesenchymal-to-epithelial transition (Figure S3E). The Serum node drives inhibition of Lox, a pro-EMT enzyme whose suppression is known to facilitate mesenchymal-to-epithelial transition during reprogramming, alongside repression of genes associated with inflammatory signaling (Ereg, Tnfrsf12a) (Figure S3G). Together, these stimulus-specific regulatory programs are individually more interpretable than a single averaged node, and broadly consistent with the known biology of the Schiebinger protocol.

## Data and Code availability

The algorithm CardamomOT, as well as the codes for generating the figures of this article, are available on https://github.com/eliasventre/CardamomOT, and the website describing the package can be found at https://cardamomot.readthedocs.io/en/latest/.

Data analyzed in this article are available at :

- Semrau et al. (2017): GEO accession GSE79578.
- Kameneva et al. (2021): GEO accession GSE147821.
- Schiebinger et al. (2019): GEO accession GSE106340 and https://singlecell.broadinstitute.org/single_cell/study/SCP295/

## Acknowledgment

We would like to especially thank Olivier Gandrillon for sharing the preprocessed dataset obtained from Kameneva et al. [39], critical reading of the manuscript and substantial help for testing and improving the Python package CardamomOT. We also thank all members of the Inria COMPO team, and of the Southrock consortium, for providing such stimulating working environment.

This work was supported by an ENS de Lyon CDSN PhD fellowship awarded to YM. The funders had no role in study design, data collection and analysis, decision to publish, or preparation of the manuscript.

## Supporting information

**Table S1:** Comparison of reference parameters and parameters inferred by CardamomOT for different simulated datasets. Results show mean *±* standard deviation across all genes.

| Parameter | FN4 |  | CN5 |  | BN8 |  | FN8 |  |
| --- | --- | --- | --- | --- | --- | --- | --- | --- |
|  | Ref | Inf | Ref | Inf | Ref | Inf | Ref | Inf |
| $k_0/d_0$ | 0 | 0.194±0.211 | 0 | 0.263±0.120 | 0 | 0.138±0.117 | 0 | 0.076±0.046 |
| $k_1/d_0$ | 10 | 8.044±0.888 | 4 | 3.315±0.401 | 8 | 5.599±1.870 | 5 | 3.474±1.445 |
| $k_{off}/s_0$ | 0.02 | 0.018±0.001 | 0.02 | 0.019±0.001 | 0.02 | 0.017±0.003 | 0.02 | 0.019±0.005 |
| % of correctly attributed modes |  | 92.9 |  | 85.2 |  | 91.3 |  | 92.6 |

**Table S2:** Related to Figure 3. Average runtime for inferring a network from datasets of 1000 cells simulated with tree-like networks for which the results of the inference are represented in Figure 3, for the six algorithms that are used in the benchmark and 10 runs each. Timings measured on a 16-GB RAM, 2.4 GHz Intel Core i5 computer.

| Runtime | PIDC | SINCERITIES | CARDAMOM | RF | **CardamomOT** | GENIE3 |
| --- | --- | --- | --- | --- | --- | --- |
| 5 genes | 0.02 s | 0.01 s | 0.10 s | 2.71 s | 4.71 s | 9.02 s |
| 10 genes | 0.04 s | 0.03 s | 0.29 s | 3.45 s | 8.92 s | 17.27 s |
| 20 genes | 0.06 s | 0.06 s | 0.72 s | 4.05 s | 17.18 s | 31.17 s |
| 50 genes | 0.18 s | 0.24 s | 2.98 s | 5.91 s | 28.57 s | 83.18 s |
| 100 genes | 0.75 s | 0.77 s | 9.14 s | 10.06 s | 55.08 s | 159.42 s |

**Table S3:** Top 10 candidate hub regulators ranked by outgoing regulatory strength Σ _*j ≠ i*_ |*θ*_*ij*_| in the GRN inferred by CardamomOT for each experimental dataset, excluding virtual stimulus nodes. Values show mean *±* standard deviation across 5 independent inference runs.

| Dataset | Rank | Gene | Out. Strength | Top 4 Outgoing Targets |
| --- | --- | --- | --- | --- |
| Semrau | 1 | Hoxb2 | 133.172 ± 8.280 | Zfp42, Pou5f1, Cd24a, Dppa2 |
| | 2 | Dnmt3a | 74.666 ± 7.691 | Nr0b1, Dppa2, Zic3, Dppa4 |
| | 3 | Lamb1 | 59.402 ± 17.010 | Sparc, Col4a2, Dab2, Gata6 |
| | 4 | Col4a2 | 44.181 ± 12.778 | Lamb1, Sparc, Gata4, Krt8 |
| | 5 | Hoxa1 | 29.536 ± 5.789 | Pou5f1, Zfp42, Hoxb2, Meis1 |
| | 6 | Esrrb | 27.715 ± 4.956 | Nr0b1, Dnmt3b, Dnmt3a, Klf4 |
| | 7 | Klf2 | 27.534 ± 6.399 | Zic2, Dnmt3b, Dppa4, Zic3 |
| | 8 | Sox2 | 23.893 ± 4.929 | Pou5f1, Esrrb, Zic3, Zfp42 |
| | 9 | Fgf4 | 22.624 ± 4.434 | Klf2, Dppa2, Nr0b1, Jarid2 |
| | 10 | Dppa2 | 21.308 ± 3.784 | Hoxb2, Zfp42, Esrrb, Klf4 |
| Kameneva | 1 | RPL30 | 152.159 ± 8.826 | RPL32, RPL10, ALDH1A1, RPL12 |
| | 2 | CHGA | 127.811 ± 15.115 | CHGB, CARTPT, SCG2, CNTN1 |
| | 3 | MEG3 | 115.906 ± 6.449 | RPL12, COL18A1, MEG8, NRXN1 |
| | 4 | POSTN | 65.639 ± 7.985 | RPL10, FAU, RPS13, RPL41 |
| | 5 | ALDH1A1 | 63.319 ± 7.361 | MAP1B, PTN, RPL32, IGFBP2 |
| | 6 | HAND2 | 61.796 ± 18.502 | PHOX2A, BASP1, UCHL1, MAP1B |
| | 7 | HAND2-AS1 | 52.155 ± 7.682 | MAP1B, UCHL1, CD24, RTN1 |
| | 8 | ATP5F1E | 49.231 ± 12.536 | RPS2, RPL34, RPL32, RPL30 |
| | 9 | RGS5 | 47.065 ± 8.048 | STMN2, RGS4, DBH, BASP1 |
| | 10 | MT-CYB | 42.587 ± 7.282 | PTPRZ1, GAL, ALDH1A1, PLP1 |
| Schiebinger | 1 | Fabp3 | 73.662 ± 9.778 | Tcl1, Mybl2, Dppa5a, Mylpf |
| | 2 | Dppa5a | 46.461 ± 5.487 | Tcl1, Cdkn2a, Lncenc1, Esrrb |
| | 3 | Ereg | 26.735 ± 5.240 | Fam20c, Myc, Ddit3, Id3 |
| | 4 | Obox6 | 23.778 ± 4.661 | Spic, Dppa4, Hesx1, Lncenc1 |
| | 5 | Txnip | 23.507 ± 4.308 | Hist1h2ap, Nme2, 2810417H13Rik, Cenpm |
| | 6 | Dmrtc2 | 22.249 ± 4.361 | Tmem35, Epcam, Id3, Cdkn2a |
| | 7 | Cdkn2a | 22.151 ± 6.199 | 2810417H13Rik, Tax1bp3, Gabarap, Fos |
| | 8 | S100a6 | 22.091 ± 7.778 | Dppa5a, Col1a2, Prss23, Lox |
| | 9 | Tnfrsf12a | 21.375 ± 5.143 | Ereg, Myc, 2810417H13Rik, Nmt1 |
| | 10 | Gm26917 | 20.426 ± 4.003 | Vps8, Vbp1, Erdr1, Scand1 |

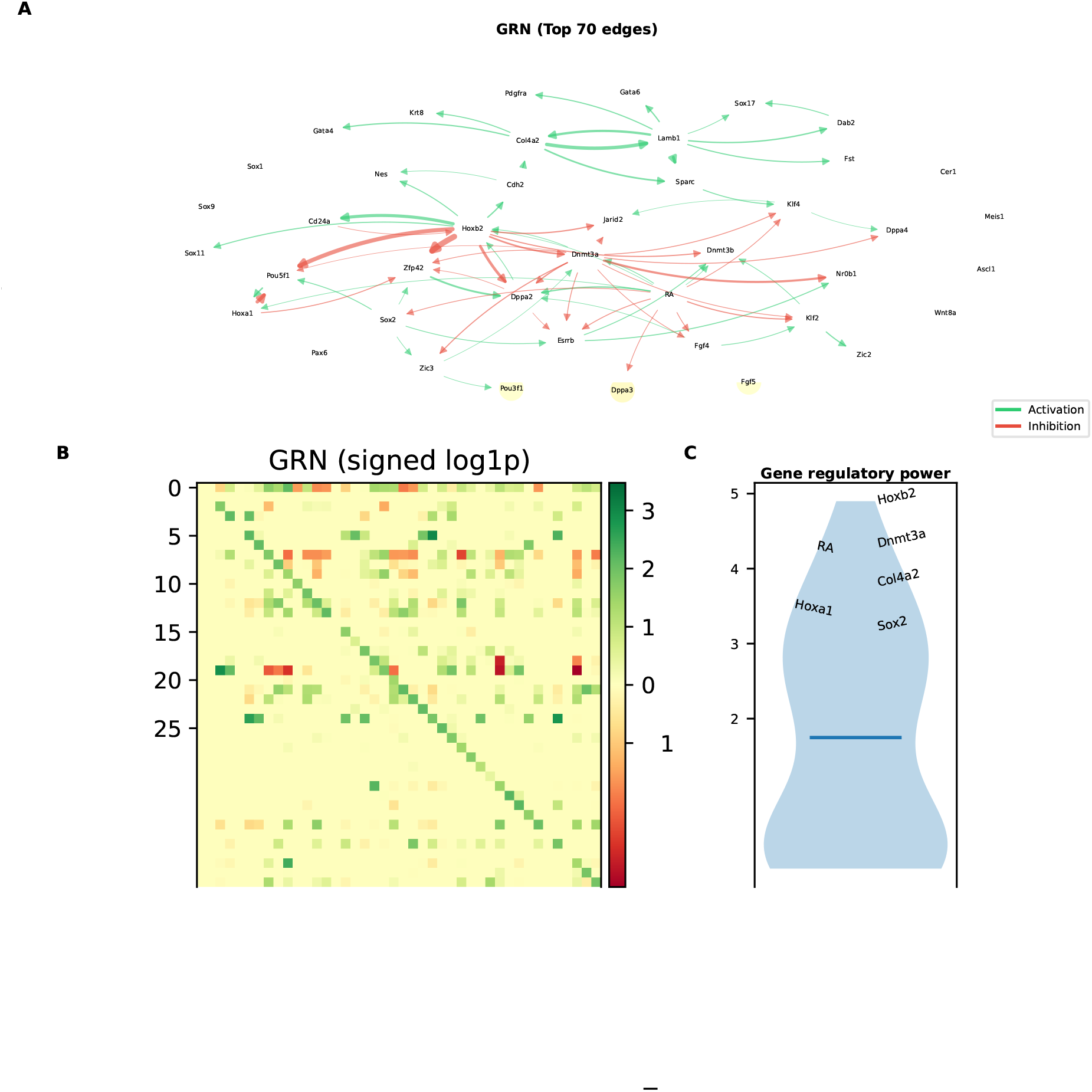

**Figure S2:**
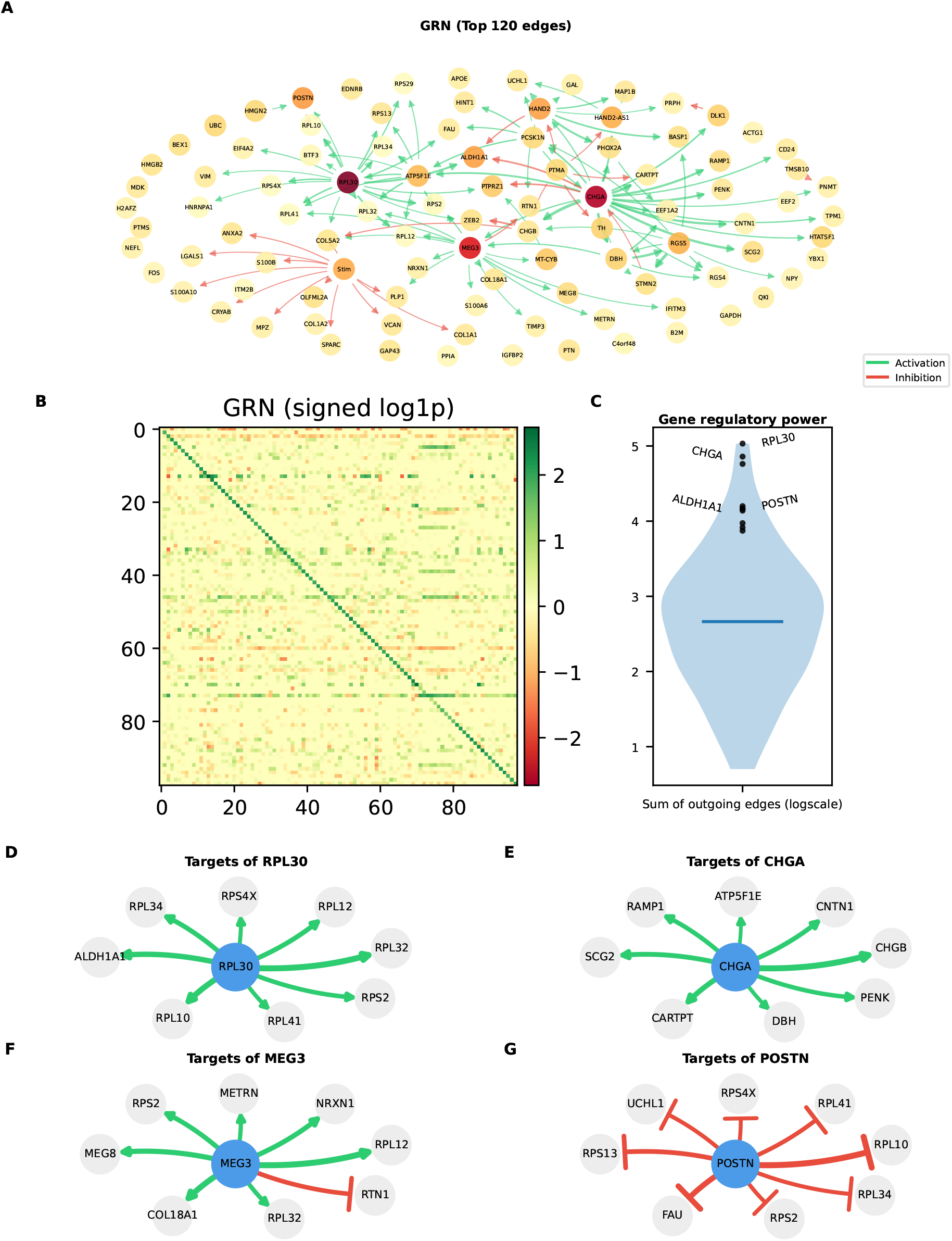
Representation of the inferred gene regulatory network for the Kameneva dataset. (A) Graph showing the top 120 edges of the GRN by absolute regulation strength, limited to a max of 5 incoming regulating edges for visualization. (B) Signed log1p transformed full GRN matrix representing the strength of all genes on each other, with lines being regulators and columns being targets. (C) Violin plot of dataset genes ranked by the sum of all their outgoing edges, with the top genes labeled. (D–G) Small GRN representation of the top regulators RPL30, CHGA, MEG3 and POSTN, showing their top 8 regulatory targets.

**Figure S3:**
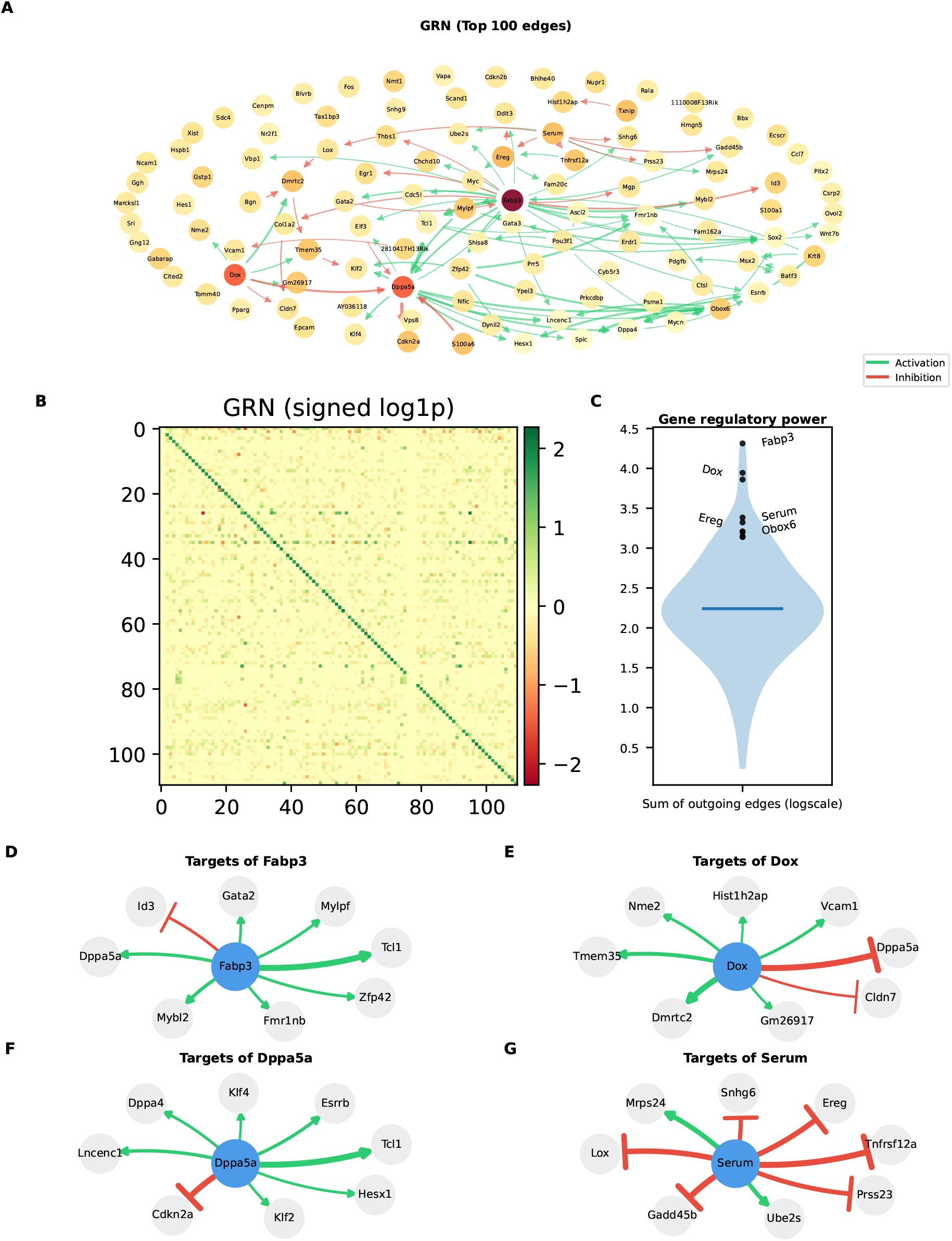
Representation of the inferred gene regulatory network for the Schiebinger dataset. (A) Graph showing the top 100 edges of the GRN by absolute regulation strength, limited to a max of 5 incoming regulating edges for visualization. (B) Signed log1p transformed full GRN matrix representing the strength of all genes on each other, with lines being regulators and columns being targets. (C) Violin plot of dataset genes ranked by the sum of all their outgoing edges, with the top genes labeled. (D–G) Small GRN representation of the top regulators Fabp3 and Dppa5a, and of the two stimulus nodes Dox and Serum, showing their top 8 regulatory targets.

**Figure S4:**
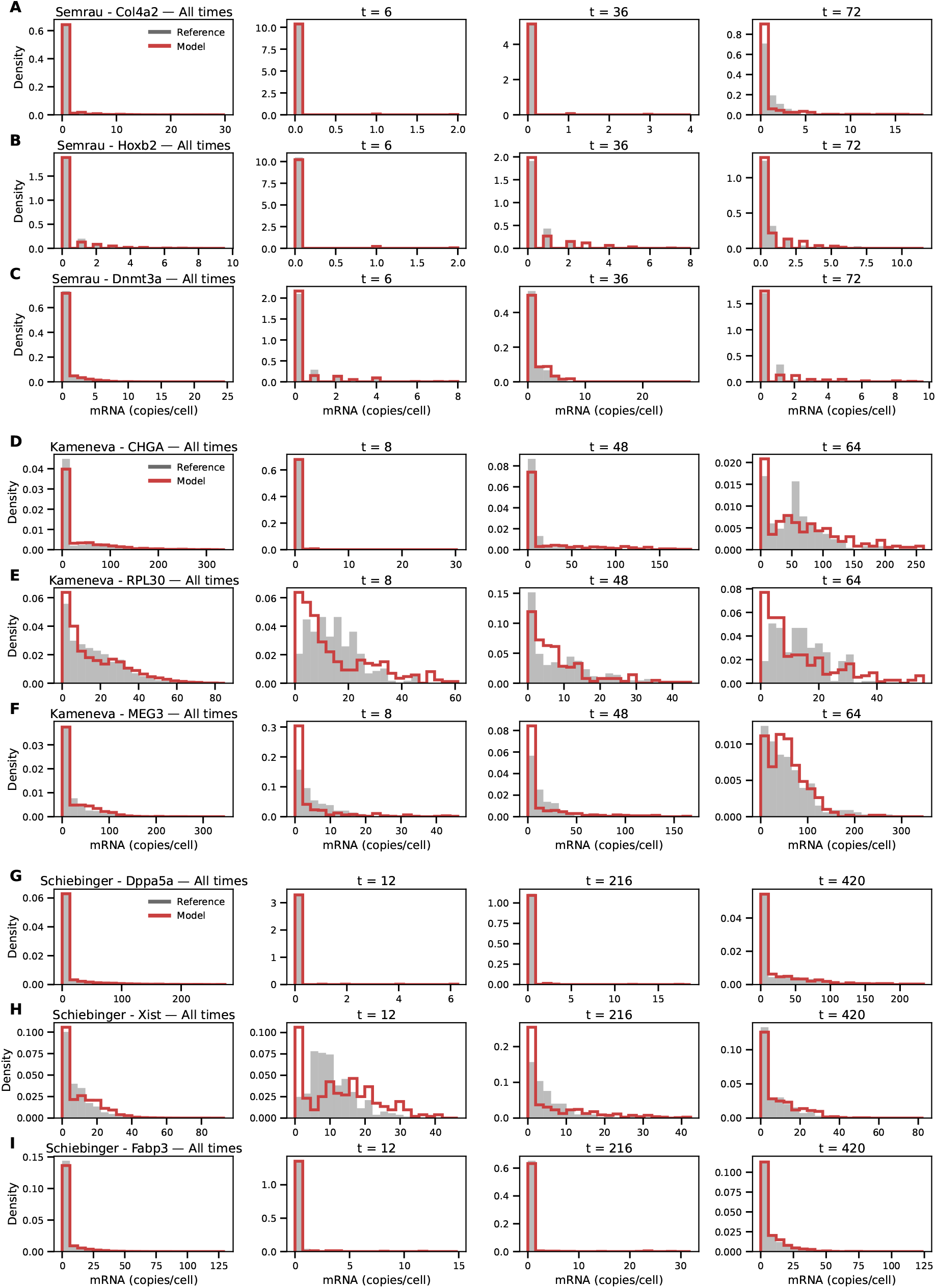
Marginal mRNA distributions for selected genes, showing (leftmost column) all timepoints joined and (remaining columns) three selected timepoints for each gene. Three of the main regulators were selected for the Semrau (A–C), Kameneva (D–F) and Schiebinger (G–I) datasets, namely *Col4a2, Hoxb2* and *Dnmt3a* for Semrau, *CHGA, RPL30* and *MEG3* for Kameneva, and *Dppa5a, Xist* and *Fabp3* for Schiebinger. Marginal distributions are shown for the reference data (gray) and the Negative Binomial mixture model (red).

**Figure S5:**
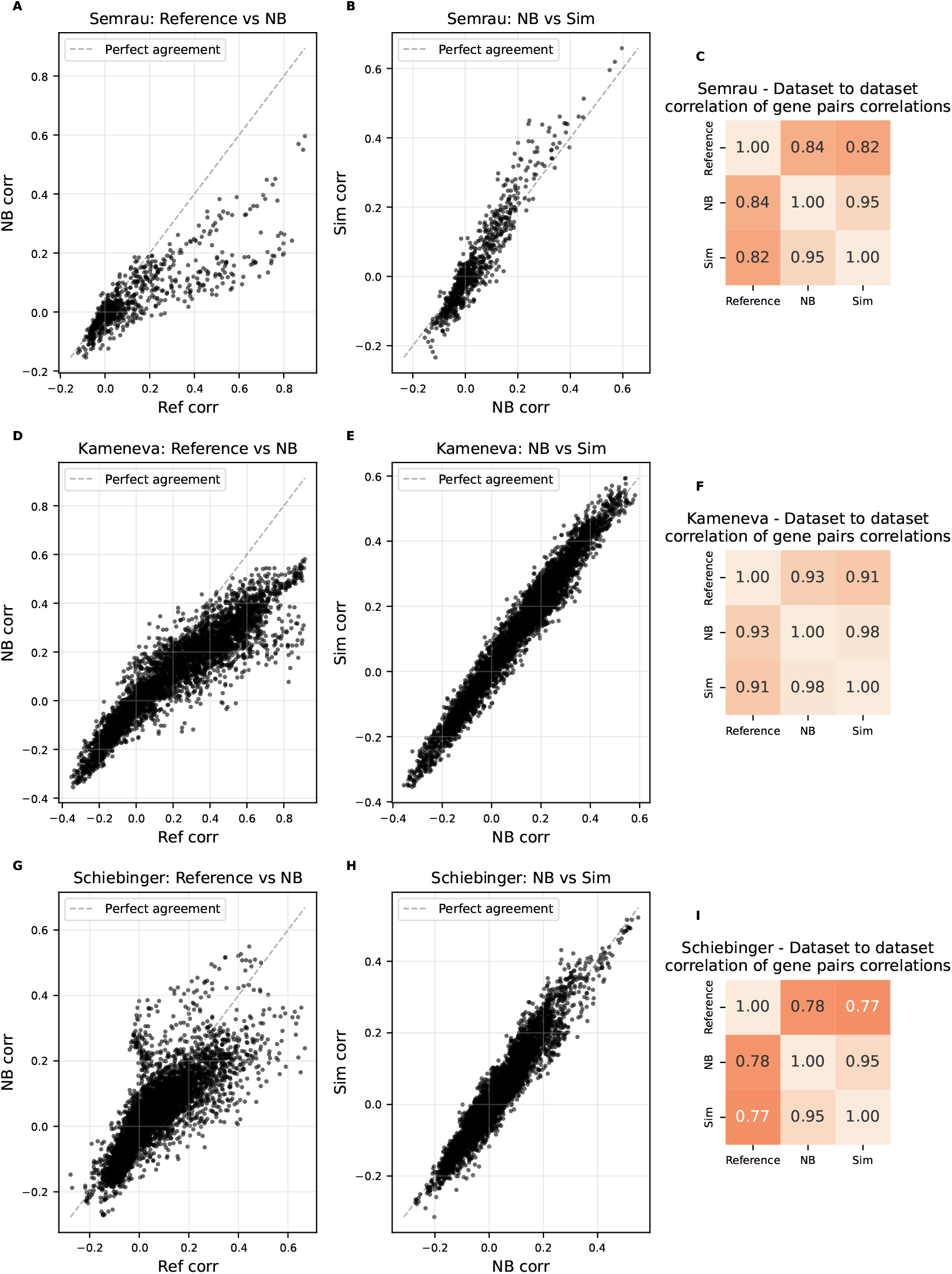
Pairwise gene expression correlation structure across reference, NB mixture and simulated datasets, for the Semrau (A–C), Kameneva (D–F) and Schiebinger (G–I) datasets. (A, D, G) Scatter plots of pairwise gene correlations computed from the reference data versus those computed from the NB mixture model, with the dashed line indicating perfect agreement. (B, E, H) Same comparison between the NB mixture model and the full generative model simulations. (C, F, I) Dataset-to-dataset correlation matrices summarizing the Pearson correlation between the vectors of all pairwise gene correlations across the three datasets (Reference, NB, Sim), providing a global measure of how well each model reproduces the correlation structure of the reference data.

**Figure S6:**
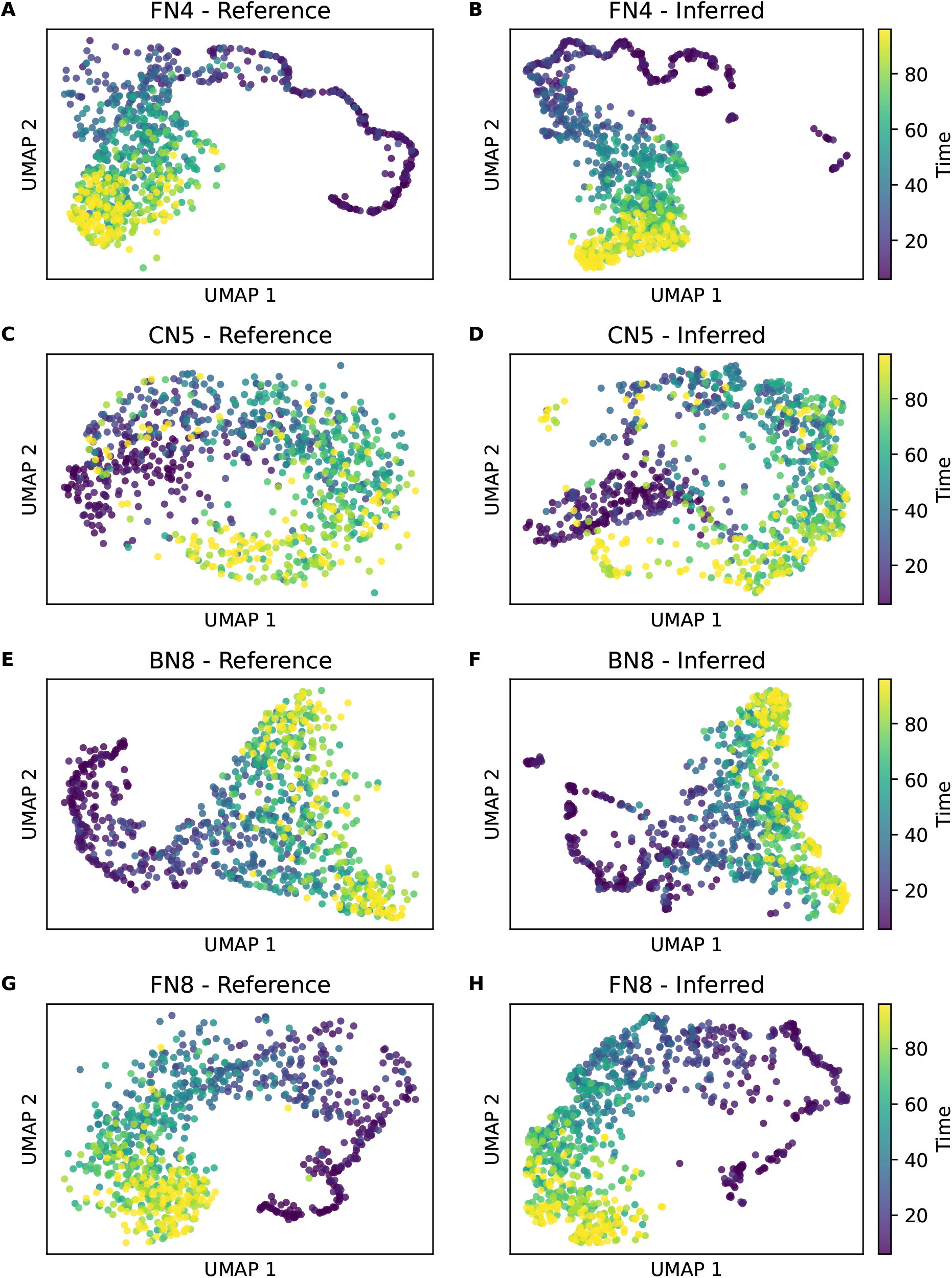
UMAPs of reference and CardamomOT inferred protein trajectories for the (A–B) FN4, (C–D) CN5, (E–F) BN8 and (G–H) FN8 datasets. (A, C, E, G) UMAPs of reference simulated protein trajectories are reproduced by (B, D, F, H) inferred protein trajectories from the reference protein-associated RNA data.

**Figure S7:**
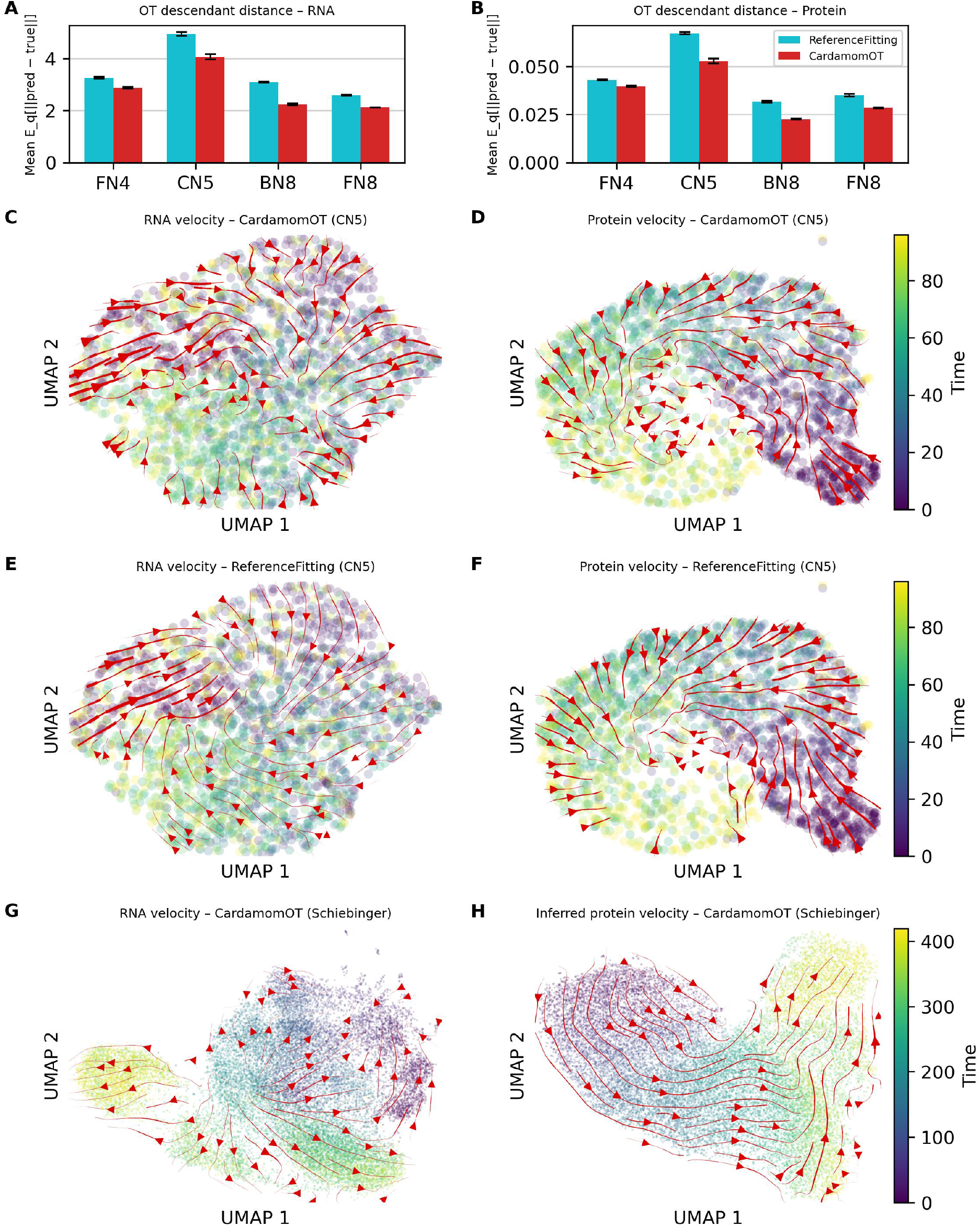
Evaluation of the cell-to-cell coupling accuracy on true trajectory data for CardamomOT and Reference Fitting. (A–B) OT descendant distance for mRNA (A) and protein (B): Bars show the mean over 5 independent simulations of true trajectories per network; error bars indicate the standard error. (C–F) UMAP embeddings of RNA (C,E) and protein (D,F) expression of CN5 benchmark cells simulated as true trajectories, with velocity stream plots derived from the inferred couplings of CardamomOT (C–D) and Reference Fitting (E–F). (G– H) UMAP of RNA (G) and inferred protein (H) data from Schiebinger et al. 2019 [15], with CardamomOT velocity stream plots derived from the inferred trajectories.

**Figure S8:**
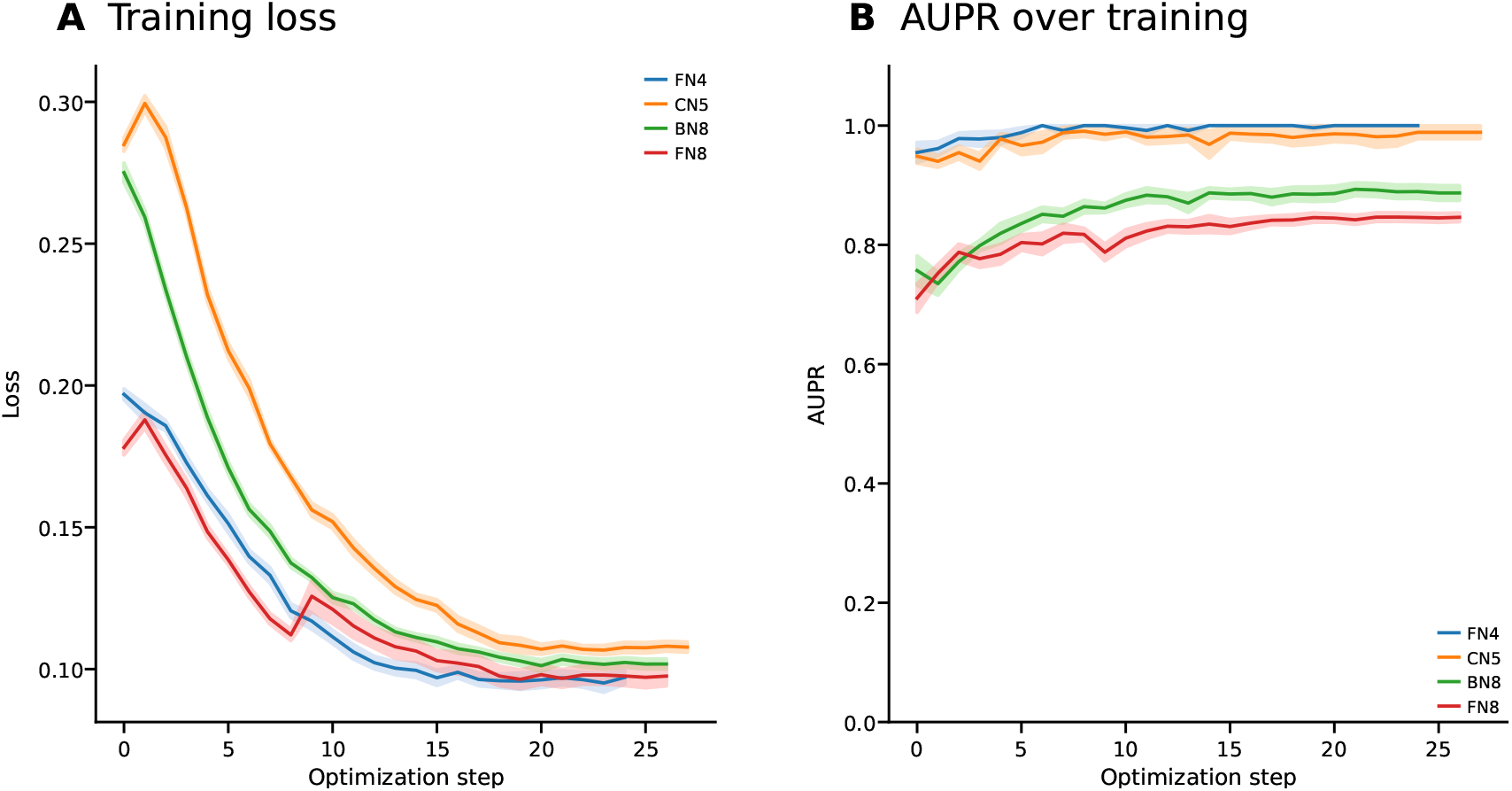
Evolution of the loss function and AUPR score during training for the networks FN4, CN5, BN8, and FN8 used in the Benchmark of Figures 3 and 4. (A) Mean training loss at each optimization step, averaged over 10 independent runs, with shaded bands indicating the uncertainty (standard error). (B) The corresponding evolution of the AUPR score over the same training steps and networks.

**Figure S9:**
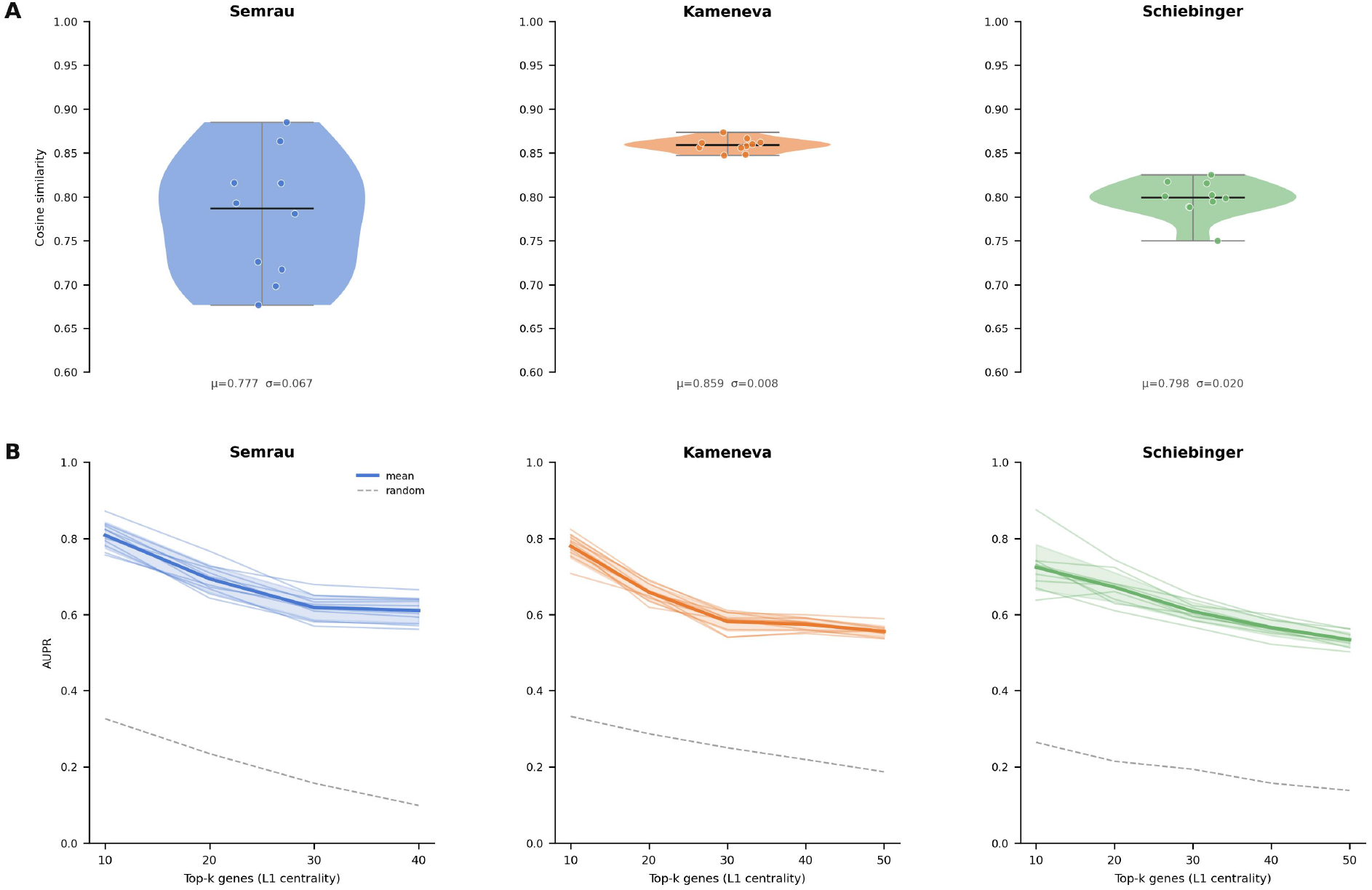
For each of the three experimental datasets (Semrau, Kameneva, Schiebinger), the inference was repeated *N* = 5 times independently with distinct random initializations. All 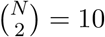 pairwise comparisons between runs are reported symmetrically, without any reference run. **(A)** Pairwise cosine similarity between GRN weight matrices across independent runs, displayed as violin plots with individual pairwise values overlaid. The mean (*µ*) and standard deviation (*σ*) are reported below each violin. **(B)** Pairwise Area Under the Precision-Recall curve (AUPR) as a function of the number of top-ranked genes considered (*k* = 10, 20, 30, 40, 50), where genes are ranked by their average L1 degree across all runs. For each pair of runs (*i, j*), the GRN from run *i* is binarized (edges with |*w*| *>* 0.5) and used as ground truth to evaluate the ranking provided by run *j*, and vice versa; the two resulting AUPR values are averaged. Thin lines represent individual pairwise comparisons; the bold line and shaded band show the mean *±*1*σ* across pairs. The dashed grey line indicates the random baseline (network sparsity at each *k*). AUPR values substantially exceed the random baseline across all datasets and values of *k*, confirming that the inferred GRN topology is reproducible across independent runs.

**Figure S10:**
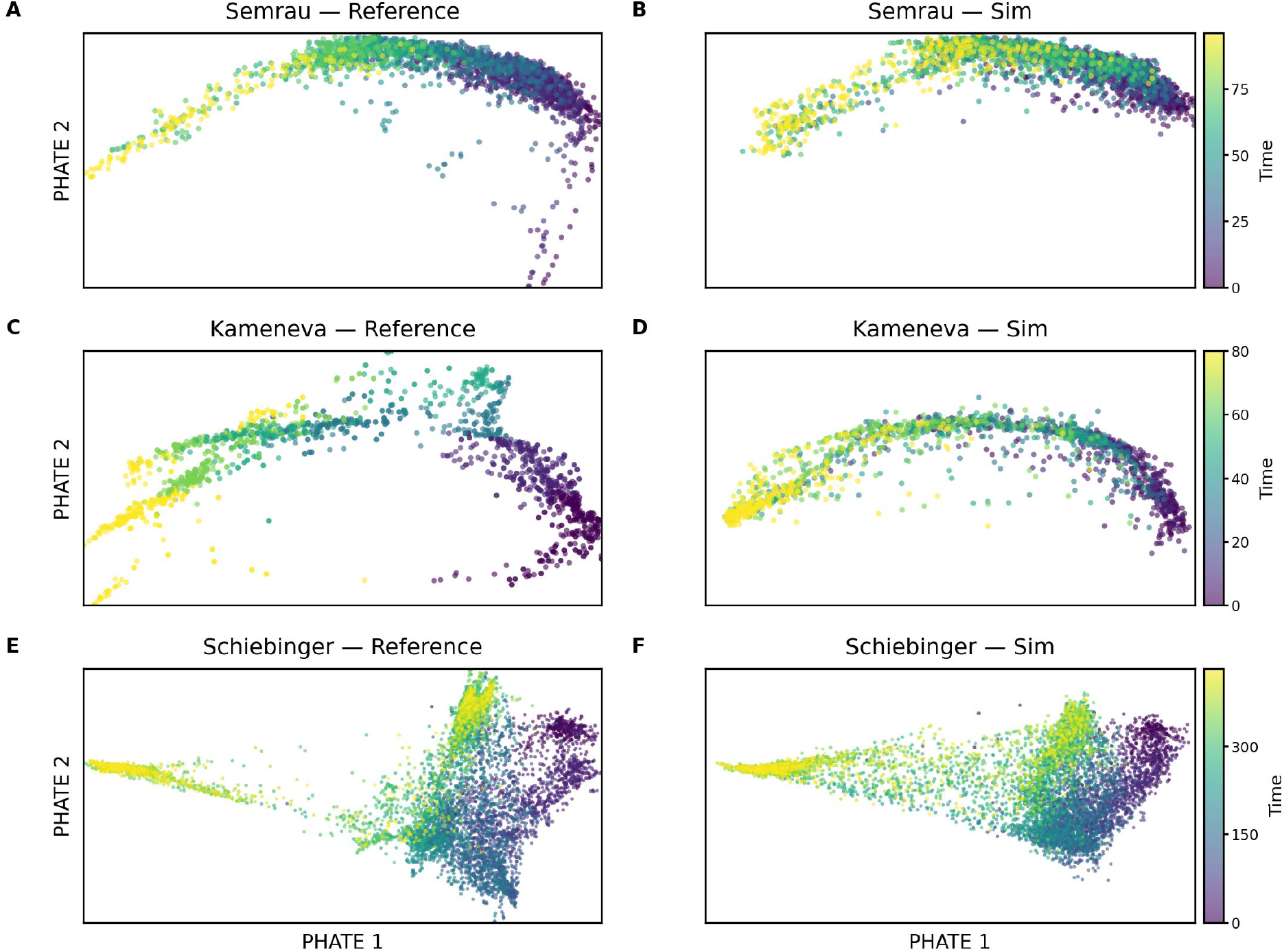
CardamomOT simulations broadly preserve the global structure and temporal ordering in trajectory-preserving PHATE embeddings. PHATE embeddings of reference (A,C,E) and simulated (B,D,F) RNA data for the 3 differentiation datasets used in the study.

**Figure S11:**
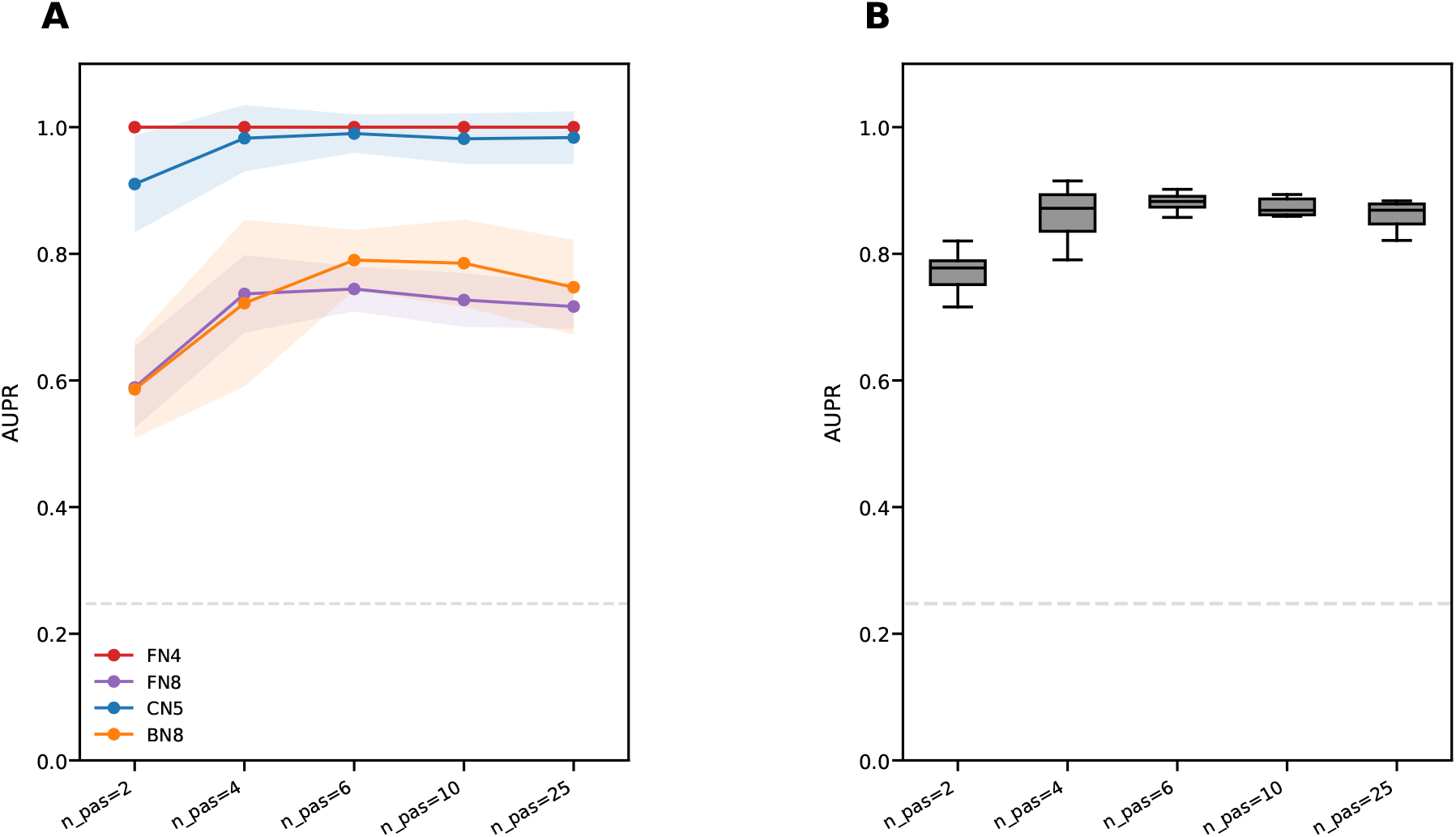
Sensitivity of GRN inference accuracy to the switching-time grid resolution *n*_pas_. (A) AUPR of CardamomOT across *n*_pas_ levels for each simulated benchmark network (FN4, FN8, CN5, BN8), shown as mean *±* SD over 10 independent simulations. (B) AUPR for the same runs averaged across the four benchmark networks. Boxplots show median, interquartile range (box) and 1.5x interquartile range (bars). The dashed gray line indicates the random baseline.

